# Generalizable virtual staining from histopathology to spatial proteomics across biological scales

**DOI:** 10.64898/2026.11.02.763811

**Authors:** Yufan Liu, Oded Falik, Zihan Zhong, Lequan Yu, Ruibang Luo, Cheng Jin, Hao Chen, Aaron T. Mayer, Alexandro E. Trevino, Peter Chou, Yue Dai, Zhenqin Wu

**Affiliations:** School of Computing and Data Science, The University of Hong Kong, Hong Kong SAR, China; Strand AI Bio Corp., San Francisco, USA; Eight-Year Medical Doctor Program, Chinese Academy of Medical Sciences & Peking Union Medical College, Beijing 100730, China; Clinical Cancer Center, National Cancer Center/National Clinical Research Center for Cancer/Cancer Hospital, Chinese Academy of Medical Sciences and Peking Union Medical College, Beijing 100021, China; Department of Computer Science and Engineering, Hong Kong University of Science and Technology, Hong Kong SAR, China; Department of Chemical and Biological Engineering, Hong Kong University of Science and Technology, Hong Kong SAR, China; Division of Life Science, Hong Kong University of Science and Technology, Hong Kong SAR, China; HKUST Shenzhen-Hong Kong Collaborative Innovation Research Institute, Futian, Shenzhen, China; State Key Laboratory of Nervous System Disorders, The Hong Kong University of Science and Technology, Hong Kong SAR, China; SIAT-HKUST Joint Laboratory for Brain Science, The Hong Kong University of Science and Technology, Hong Kong SAR, China; Enable Medicine, Menlo Park, CA, USA; School of Biomedical Engineering, The University of Hong Kong, Hong Kong SAR, China

**Author notes:** These authors contributed equally to this work (Yufan Liu, Oded Falik).

## Abstract

Large histology archives capture morphological correlates of molecular tissue states but lack direct spatial protein measurements. In this study, we show that routine hematoxylin and eosin (H&E) histology can provide a queryable, morphology-based representation of spatial protein expression that remains informative from pixels to patients. To achieve this, we developed QUEST, a model framework that represents protein markers as semantic queries for flexible multiplex immunofluorescence (MIF) prediction. Trained on a heterogeneous atlas of paired H&E and MIF images comprising over 10,000 tissue regions, QUEST captured morphological patterns associated with protein markers and reconstructed their expression intensities across diverse tissue types and cohorts. Its utility extended beyond image reconstruction and preserved biologically meaningful information across multiple scales: virtual staining preserved protein marker expression at the cellular and tissue scales, supported cell phenotyping, cellular neighborhood analysis, and characterization of spatial tissue organization, retained clinically relevant signals for patient stratification, and enabled H&E-based queries to retrieve MIF references with similar molecular profiles. These findings position virtual staining not simply as image generation but as a flexible interface between routine histology and spatial proteomics, expanding the analytical value of existing H&E archives.

## Introduction

Hematoxylin and eosin (H&E) staining-based histology, which provides detailed views of tissue architecture and cellular morphology, is foundational to the diagnosis of diverse diseases. Beneath these morphological patterns lies a layer of molecular organization that shapes tissue behavior, including different immune cell lineages, programs related to activation, immune checkpoints, exhaustion, and stromal function. Although some molecular programs correlate with morphological patterns, they are not directly measured by H&E staining. Multiplex tissue imaging complements morphology by profiling protein expression *in situ* and revealing recurrent cellular neighborhoods, tumor-host interactions, and diverse niches linked to disease progression and treatment response (Hickey et al. 2023; Keren et al. 2018; Schürch et al. 2020; Asrir et al. 2022; J.-R. Lin et al. 2023).

Generating protein expression maps remains expensive and resource-intensive, limiting their applications in large cohorts. By contrast, H&E staining is routinely performed and therefore widely available in both clinical practice and tissue archives. Importantly, previous studies have demonstrated that deep learning can identify morphological patterns in H&E-stained images that are associated with genetic alterations and molecular phenotypes (Coudray et al. 2018; Kather et al. 2019, 2020). These findings suggest that some aspects of the underlying molecular landscape are reflected in tissue morphology. Together, the widespread availability of H&E-stained images and growing evidence that tissue morphology contains correlates of the underlying molecular state have motivated efforts to predict spatial protein-expression maps directly from histology.

Recent efforts have demonstrated the feasibility of predicting multiplex immunofluorescence (MIF) images from H&E images across increasingly diverse cohorts (Wu et al. 2025; Valanarasu et al. 2026; Balezo et al. 2026; Li et al. 2026). However, most virtual MIF staining methods remain formulated as image translation to a predefined marker panel. This design limits their reuse across studies with different marker panels and leaves generalization to new markers largely unresolved. Robust generalization across datasets also remains challenging, as tissue type, disease context, staining platform, and marker prevalence can influence the relationship between H&E-based morphology and protein expression. Beyond these limitations, although several studies have explored the downstream biological and clinical applications of virtual MIF staining, such capabilities have not been systematically benchmarked across models, cohorts, and tasks. Further, the potential of virtual MIF staining to enable cross-modal applications that connect H&E-based histology with archives of existing spatial proteomics data also remains underexplored.

In this study, we developed QUEST (**QUE**ry-based virtual **ST**aining) to address these gaps. Rather than assigning each marker to a fixed output panel, QUEST represents requested markers as queries that interact with H&E features and share a common spatial readout. This formulation allows the requested panel to be specified at inference and creates a natural route for transfer to markers not observed during training. QUEST was trained and evaluated on a large, heterogeneous atlas of paired H&E and multiplex immunofluorescence images comprising over 2.4 million patches spanning diverse tissues and marker panels.

We systematically evaluated the predictive fidelity and biological utility of QUEST across held-out and external cohorts at multiple scales, ranging from pixels and cells to tissue regions and patients. Beyond image reconstruction, we examined whether virtual MIF preserved biologically meaningful information for cell phenotyping, spatial tissue characterization, and downstream clinical analyses. We further evaluated the utility of virtual MIF for cross-modal applications. Our results show that virtual MIF preserves biologically and clinically relevant information across scales, from cellular phenotypes and tissue organization to patient-level endpoints. Together, we envision QUEST transforming virtual MIF staining from fixed-panel image translation into a queryable bridge between routine H&E and spatial proteomics, opening a novel path to molecularly informed analysis of the vast H&E histology collections already available in research and clinical practice.

## Results

### Heterogeneous H&E-to-MIF datasets for model development and benchmarking

To support the development of H&E-to-MIF virtual staining models across diverse tissue contexts and marker panels, we assembled a large, heterogeneous collection of spatially co-registered paired H&E and MIF images (**Figure 1a**). Whole-slide images and tissue microarrays were cropped into non-overlapping image patches for model training and held-out evaluation (Vorontsov et al. 2024; R. J. Chen et al. 2024). We curated data spanning diverse tissue types and imaging markers. The collection for model development and evaluation comprised 12,880 tissue regions from 10,891 patients, from which more than 2.4 million image patches were curated, and 93% of the regions were obtained from cancer tissues. In total, the collection contained 129 distinct biomarkers, with a detailed list provided in **Supplementary Data 1**. Among these, 29 broadly expressed markers were assayed across all tissue types (**Figure 1b, Supplementary Figure 1a**). Marker panels varied across regions, with a median panel size of 53 (**Supplementary Figure 1b**). Colon and rectum represented the largest tissue type category, comprising more than 1,500 regions, whereas stomach, breast, lung, reproductive system, and pancreas each contributed more than 1,000 regions. A held-out collection containing 1,812 regions from 1,769 distinct patients was used for testing and downstream evaluations (**Figure 1c**), based on which we curated test datasets including CRC (colorectal cancer)-stain, Breast-stain, Melanoma-stain, Lung-stain, and a Mixed 10K dataset that comprises 10,000 patches randomly sampled from all tissue types (**Methods**). In addition, we used multiple external evaluation cohorts. The HEMIT (Bian et al. 2024), ORION-CRC (J.-R. Lin et al. 2023), and HuBMAP-intestine (Hickey et al. 2023; HuBMAP Consortium 2019) datasets were used to assess virtual MIF staining fidelity across different platforms. H&E images and clinical annotations from The Cancer Genome Atlas (TCGA, Cancer Genome Atlas Research Network et al. 2013) were employed to assess downstream clinical applications.

**Figure 1:**
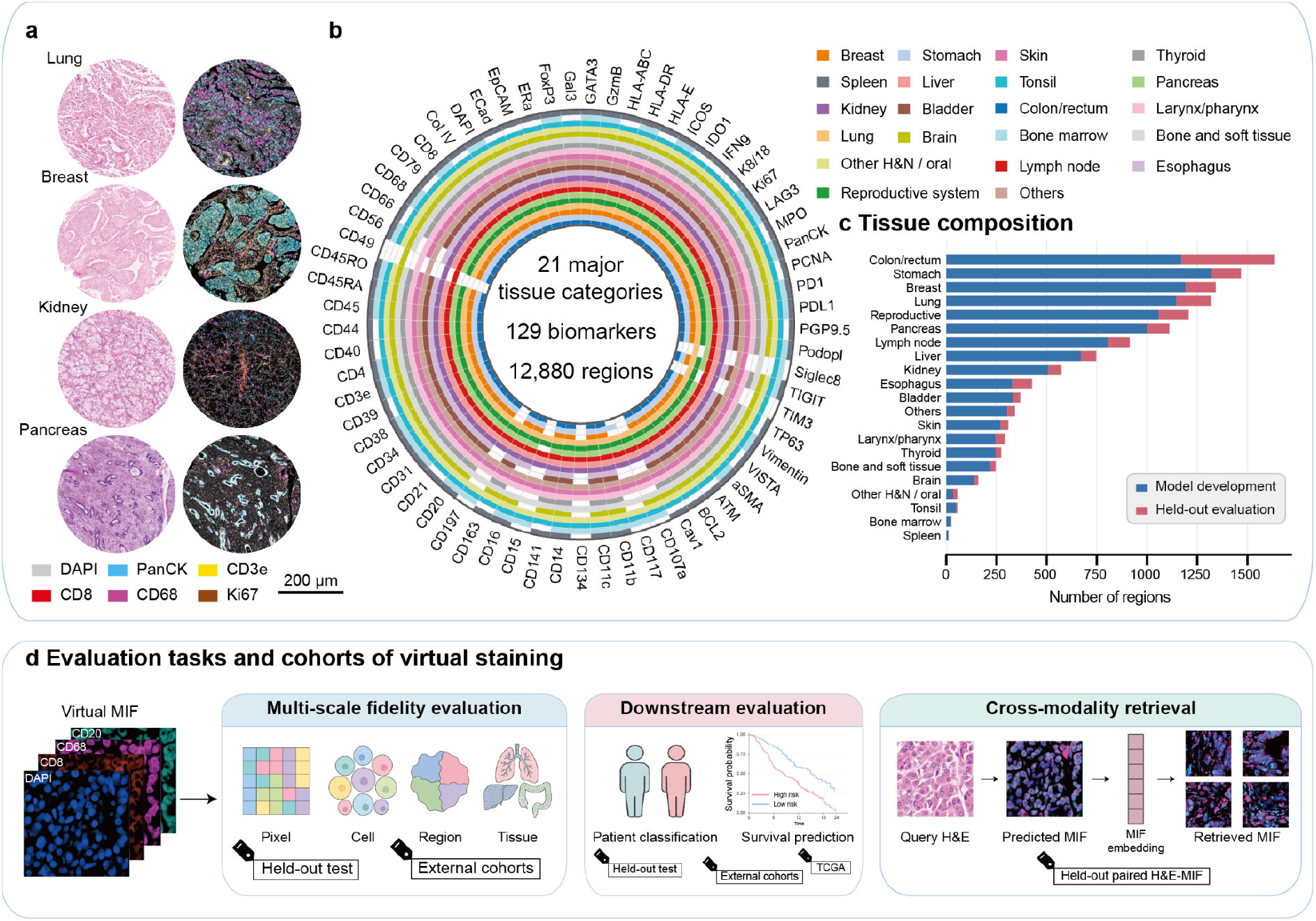
Datasets for model development and evaluation. **a**, Representative paired H&E and multiplex immunofluorescence (MIF) images from lung, breast, kidney, and pancreas tissue microarray cores. **b**, Coverage of selected representative biomarkers across tissue types. The complete dataset comprises 129 biomarkers, 21 major tissue categories, and 12,880 tissue regions (**Methods**). **c**, Distribution of tissue regions across model-development and held-out evaluation sets, stratified by tissue type. **d**, Overview of the evaluation framework. Virtual MIF fidelity was assessed at the pixel, cell, region, and tissue scales using held-out data and external cohorts. Downstream utility was evaluated through patient classification and survival prediction using held-out, external, and TCGA cohorts. Cross-modality retrieval was evaluated on paired held-out H&E–MIF data by retrieving MIF references from H&E-derived virtual MIF representations.

We organized the evaluation into three task categories (**Figure 1d**). First, multi-scale fidelity assessment quantified agreement between virtual and real MIF signals at the pixel, cell, region, and tissue levels using held-out and external datasets. Second, downstream evaluations examined whether virtual MIF inferred from H&E retained clinically informative signals for patient stratification and survival prediction. Third, cross-modality retrieval assessed whether virtual MIF can link an H&E query image to galleries of real MIF images. Further details of the held-out test datasets, external evaluation cohorts, and task-specific protocols are provided in **Methods**.

## Model development of QUEST

The substantial variation in protein marker composition across cohorts and studies presents a fundamental challenge for existing H&E-to-MIF virtual staining models. We developed models for flexible prediction of spatial proteomics maps from H&E images, named QUEST (**QUE**ry-based virtual **ST**aining), a dynamic, query-based architecture designed specifically for virtual MIF staining.

QUEST receives an H&E image together with a specified set of marker queries and generates one spatial intensity map for each requested marker (**Supplementary Figure 2a**). Three model variants of QUEST were developed: QUEST-semantic and QUEST-ID used the pretrained pathology foundation model UNI2-h (R. J. Chen et al. 2024) as their feature backbone. QUEST-Eva leveraged Eva (Yufan Liu et al. 2025), a masked autoencoder framework based on the Vision Transformer (ViT) model pretrained jointly on spatial proteomics and matched histopathology, as its backbone. Briefly, marker queries interact with the image representations through attention-based mechanisms, and a readout decoder uses the integrated embeddings to generate the corresponding virtual MIF maps. To capture the heterogeneous marker panels, QUEST-ID uses learned marker-specific embeddings and is therefore restricted to markers encountered during training. QUEST-semantic leverages marker representations defined from fixed text-derived embeddings within a shared semantic space, providing a route to predicting markers absent from the training vocabulary (**Methods**).

### Pixel-level virtual MIF staining across heterogeneous datasets

We assessed virtual MIF staining fidelity across held-out and external evaluation datasets spanning diverse tissues and marker panels. We compared our models against Eva (Yufan Liu et al. 2025), GigaTIME (Valanarasu et al. 2026), GigaTIME-flash (Usuyama et al. 2026),ROSIE (Wu et al. 2025), and MIPHEI-ViT (Balezo et al. 2026). Pixel-level agreement between virtual and real MIF images was assessed using the Pearson correlation coefficient (PCC) and Spearman’s rank correlation coefficient (Spearman’s ρ) at the patch level, calculated both at the original pixel resolution and after local spatial aggregation into grids (**Methods**).

QUEST models achieved higher pixel-level PCC and Spearman’s ρ than previous methods in most datasets, and their performance advantages were retained after grid-level aggregation (**Figure 2a**). Grid-level correlations were generally higher than their pixel-level counterparts, consistent with local spatial averaging reducing sensitivity to minor pixel-level misalignment and high frequency intensity variation. As existing methods were developed using different marker panels, we repeated the analysis using the intersection marker sets for each benchmarked method. Across comparisons, QUEST models generally achieved higher overall PCC **(Supplementary Figure 3a**). Notably, ORION-CRC was used to train MIPHEI-ViT (Balezo et al. 2026).

**Figure 2:**
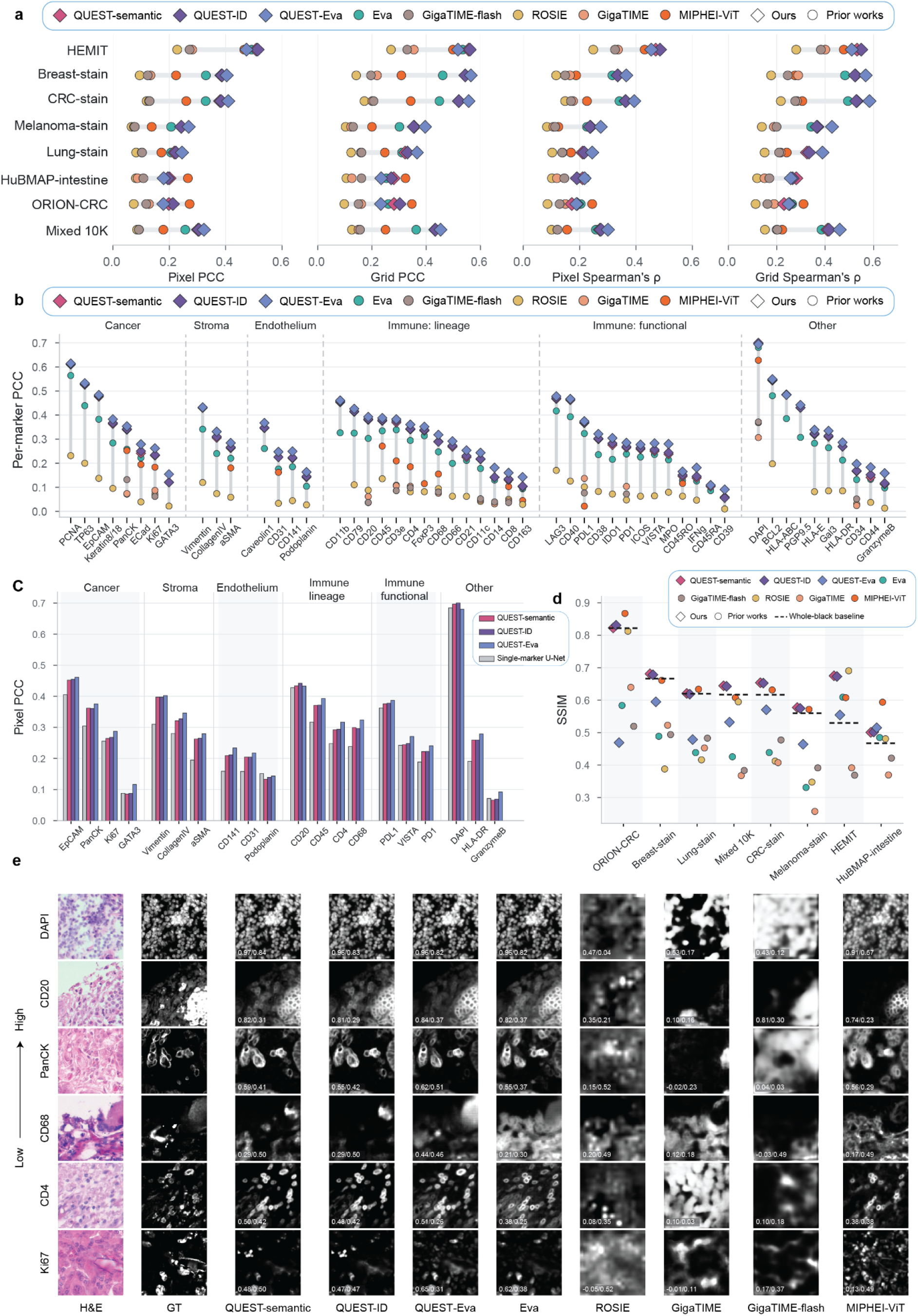
Image generation fidelity evaluation across datasets and biomarkers. **a**, Pearson correlation coefficient (PCC) and Spearman’s rank correlation coefficient (Spearman’s ρ), evaluated at the pixel-level and grid-level (14×14) aggregation across external datasets and held-out test sets (**Methods**). **b**, Per-marker pixel-level PCC on the Mixed 10K dataset, with markers grouped by biological compartment or function. **c**, Marker-wise predictability assessed using independently trained single-marker U-Nets and compared with QUEST-semantic, QUEST-ID and QUEST-Eva for representative markers. **d**, Structural similarity index measure (SSIM) across datasets. Black dashed segments indicate the SSIM achieved by all-black predictions for the corresponding datasets. **e**, Visual examples of patch-level H&E inputs, real MIF references, and virtual staining predictions from different models for markers spanning a range of prediction difficulty. Values overlaid on each prediction report PCC/SSIM.

Prediction accuracy varied considerably across markers (**Figure 2b**). QUEST models achieved strong per-marker PCC across cancer, stromal, endothelial, immune-lineage, and immune-functional groups, although absolute performance remained dependent on the abundance, morphological correlates, and predictability of each marker. Markers associated with visually distinct cellular or tissue structures generally achieved higher correlations, including DAPI, PCNA, TP63, EpCAM, and Vimentin. By contrast, sparse immune and functional markers, such as CD8, CD163, and CD39, remained more difficult to infer from H&E morphology. To this end, we further compared QUEST models with independently trained single-marker U-Net models (**Methods**, Ronneberger et al. 2015). Independently trained single-marker models provided an empirical estimate of marker-level predictability. QUEST models generally matched or exceeded the specialized single-marker models (**Figure 2c**). Our results indicate that some markers have limited predictability from H&E morphology rather than being limited by our modeling architectures.

MIF images are often sparse, with marker-positive signals occupying only a small fraction of the image. Consequently, the structural similarity index measure (SSIM, Wang et al. 2004), a different assessment from the correlation-based metrics, may be biased (**Figure 2d**). An all-black prediction containing no marker signal achieved SSIM values approaching or exceeding those of trained virtual MIF staining models, because large background regions dominated the metric even when biologically relevant signals were missed. We therefore complemented the evaluations with foreground-centric analyses that assessed both intensity agreement within positive signals and discrimination between foreground and background.

We defined foreground and background using Otsu thresholding (**Methods**, Otsu 1979). Foreground PCC, calculated within marker-positive pixels of the real MIF image, showed that QUEST models outperformed the corresponding baselines in most datasets (**Supplementary Figure 3b**). Pixel-wise scatter plots further showed closer agreement between predicted and measured signal intensities (**Supplementary Figure 3c**). We additionally evaluated foreground–background discrimination using F1 score and AUPRC, which capture both missed foreground signals and false positive predictions in background regions. QUEST models generally outperformed other virtual MIF staining methods on both metrics (**Supplementary Figure 4a–b**), and their predicted foreground masks more closely reproduced the spatial extent and distribution of marker-positive regions (**Supplementary Figure 4c**).

Together, these complementary metrics assessed the overall agreement, foreground-centric correlation, and foreground-background discrimination of virtual MIF stains. Visual examples further showed that QUEST models recovered spatial patterns of different markers with high fidelity (**Figure 2e**).

### Pan-cancer training improves tissue-specific predictions

To evaluate the benefits of pan-cancer training, we compared the pan-cancer QUEST model with identically configured models trained exclusively on colon or lung tissues using the same QUEST-semantic architecture. On tissue-specific marker panels, pan-cancer training increased macro PCC from 0.352 to 0.383 in colon and from 0.183 to 0.228 in lung. **Supplementary Figure 5a** shows the corresponding per-marker changes (ΔPCC). Among the 57 markers shared between the two tissues, the per-marker gains were positively related (PCC = 0.695), indicating that the benefits of pan-cancer training were reproducible across tissue contexts. The largest gains occurred for vascular and stromal markers, including CD34, CD31, and aSMA, whose associated morphological structures are broadly shared across tissues.

We next tested whether supervision for a marker in other tissues could generalize to its prediction in a target tissue where the marker was not observed. We selected five markers measured in both the colon and lung panels, including EpCAM, BCL2, PDL1, Ki67, and FOXP3, and constructed controlled target-tissue marker holdouts. For each tissue-specific model, these markers were removed from all training data and therefore evaluated as zero-shot markers. For the corresponding pan-cancer model, the same markers remained excluded from the target tissue but were supervised in other cancer types (**Methods**). Supervision from other tissues significantly improved all five markers in colon and three of five markers in lung, whereas Ki67 and FOXP3 in lung showed no detectable difference (**Supplementary Figure 5b)**.

### Virtual MIF preserves region-level fidelity and supports cell-level profiling

Having evaluated pixel-level fidelity of virtual MIF, we next evaluated the fidelity of virtual MIF staining across diverse tissue types using held-out test data (**Methods**). We summarized the correlations between virtual and real MIF signals across 21 tissue categories (**Figure 3a, Methods**). QUEST-Eva achieved the highest PCC of 0.275 across tissue types, followed by QUEST-ID and QUEST-semantic, outperforming existing virtual MIF staining methods (0.079–0.218). Performance varied across tissue types, with higher average correlations in colon and rectum, breast and stomach, and lower correlations in skin and bone marrow. We further demonstrated virtual MIF staining on whole slide images (WSI) from the held-out dataset and TCGA (**Supplementary Figure 6a-b**).

**Figure 3:**
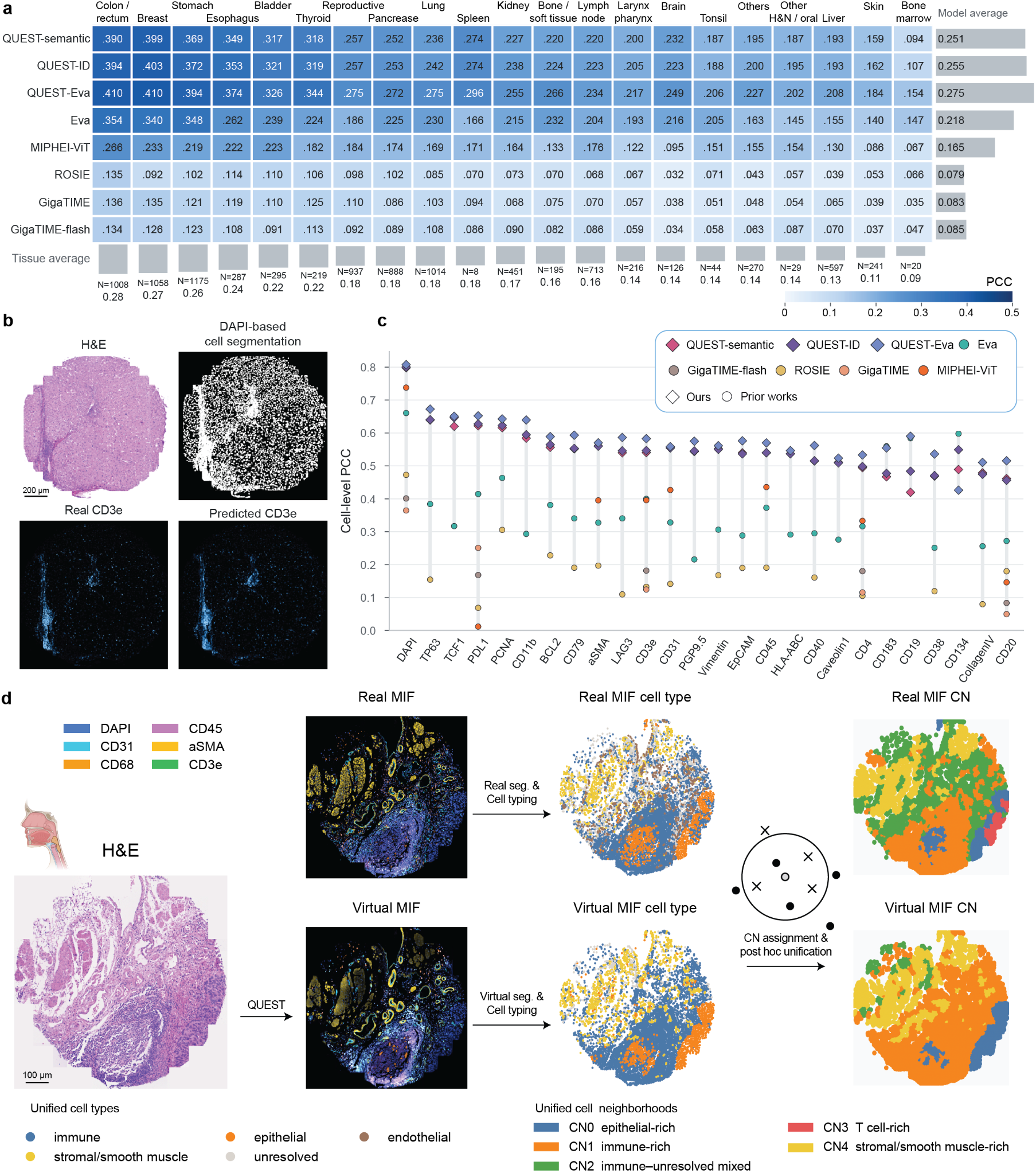
Multi-scale fidelity of virtual MIF across tissues, cells, and cellular neighborhoods. **a**, Region-level PCC between predicted and real MIF signals, stratified by tissue type and model. Cell values and color intensity indicate PCC. Gray bars on the right show model averages across tissues, whereas those at the bottom show tissue averages across models. N indicates the number of training regions for each tissue type. **b**, The input H&E image, DAPI-based cell segmentation, and real and predicted CD3e signals at cellular resolution are shown for a tissue region. **c**, Per-marker PCC between predicted and measured continuous cell-level intensities. **d**, End-to-end cellular neighborhood (CN) annotation workflow illustrated in an esophagus region. The real MIF reference used cells segmented from real DAPI and phenotyped using measured marker expression. For virtual MIF, cells were independently segmented from virtual DAPI and phenotyped using virtual marker expression. For each cell, the composition of its local neighborhood was defined from its 10 nearest spatial neighbors, and the resulting cell type composition was used to assign the cell to a CN. After independent CN construction, cell types and CNs from the real and virtual MIF were mapped post hoc to unified biological annotations for visualization and biological comparison (**Methods**). Final CN maps were visualized as Voronoi diagrams generated from cell centroids. All virtual MIF visualization examples shown in **b** and **d** are based on QUEST-semantic predictions.

We next evaluated whether virtual MIF preserved marker expression at cellular resolution. Cells were identified by a DAPI-based segmentation method (Greenwald et al. 2022), with measured and predicted marker intensities aggregated within each cell (**Figure 3b, Methods**). For each region and marker, cell-level PCC was calculated across all segmented cells (**Methods**). QUEST models formed the leading group across most markers, with QUEST-Eva achieving the highest PCC for the majority of markers (**Figure 3c**).

Many downstream analyses use binarized marker expression to characterize cellular populations (Berry et al. 2021). Thus, we additionally evaluated whether the models could distinguish marker-positive from marker-negative cells (**Methods**). QUEST models achieved consistently high cell-level AUROC across the evaluated markers and generally outperformed existing methods (**Supplementary Figure 7a**). We showed that QUEST models closely reproduced the distributions of marker-positive cells across markers with different abundance levels and spatial patterns (**Supplementary Figure 7b**). We additionally assessed whether virtual DAPI staining could support cell segmentation. Segmentation based on virtual DAPI outperformed segmentation based on H&E-derived hematoxylin, with QUEST models showing the strongest performance among the evaluated methods and producing cell instances that closely matched those segmented from measured DAPI (**Supplementary Figure 8a-b, Methods**).

We then explored whether expression profiles derived from virtual MIF could support cell phenotyping on the external PathoCell (Lüscher et al. 2026; Schürch et al. 2020) cohort. In this task, the same cell segmentation masks derived from real DAPI stains were used, and each model was evaluated using its maximum available marker panel coverage (**Methods**). On the fine-grained 15-class annotations, QUEST variants led (F1 0.246 for QUEST-semantic and 0.245 for QUEST-ID), followed by MIPHEI-ViT at 0.225. We also defined a coarse 6-class annotation of cell types (**Methods**), where similar results were observed (**Supplementary Figure 9**).

We next investigated whether virtual MIF could independently recover cellular neighborhood (CN) organization from virtual stains alone (**Methods**). We first evaluated the complete virtual workflow, in which cells were segmented from virtual DAPI, cell types were independently clustered and annotated from virtual marker expression, and CNs were subsequently defined and annotated from local virtual cell-type compositions. In an example of the esophagus region, the independently derived virtual cell types and CNs reproduced the dominant cellular organization and major CN spatial patterns observed in the real MIF reference after post hoc mapping to a unified biological annotation vocabulary (**Figure 3d, Supplementary Data 2-3**). We additionally performed a supervised comparison using the same real DAPI-derived cell masks and fixed real MIF-derived CN labels for all methods (**Methods**). Virtual MIF-based prediction generally outperformed direct prediction from H&E, with QUEST achieving the highest CN accuracy among the evaluated virtual staining methods in both esophagus and lung tissues (**Supplementary Figure 10**).

Beyond cellular neighborhood annotation, we examined whether virtual MIF staining preserved biologically relevant spatial organization of cells. Blood and lymphatic vessels provide key routes for T-cell entry, positioning, and trafficking in tissues (Asrir et al. 2022). Here, we used virtual MIF stains to quantify the distance from each T cell to its nearest vascular or lymphatic cell. QUEST models broadly recapitulated the corresponding ground-truth distance distributions (**Supplementary Figure 11a**). At the niche level, local cell type entropy estimated from virtual MIF stains showed moderate agreement with the real MIF reference; spatially, low- and high-entropy niches distinguished compositionally homogeneous regions from heterogeneous interfacial and immune activity sites (**Supplementary Figure 11b**).

### Virtual MIF recapitulates the clinical utility of real MIF and complements H&E-based prediction

Beyond image-level fidelity and cell-level profiling, we investigated whether virtual MIF could provide useful or complementary information for patient-level clinical predictions. First, we tested whether virtual MIF retained the downstream predictive utility of real MIF. In the held-out test cohorts with paired H&E and real MIF, we compared the downstream utility of virtual and real MIF using the same marker panels on 4 distinct clinical tasks. QUEST-based virtual MIF staining achieved performance comparable to real MIF (**Figure 4a**). For melanoma stage classification, QUEST-ID and QUEST-semantic achieved the highest AUROCs of 0.817 and 0.801, as compared to the real MIF reference of 0.780. For lung cancer stage classification, QUEST models led with AUROCs of 0.714–0.754. For CRC metastatic survival modeling, real MIF remained the strongest modality with a C-index of 0.758, followed by QUEST-ID and QUEST-semantic at 0.725 and 0.717, respectively, whereas existing methods achieved 0.515–0.640. For breast cancer grade classification, real MIF achieved the highest AUROC of 0.719, while QUEST models remained the leading virtual MIF staining methods at 0.667–0.689.

**Figure 4:**
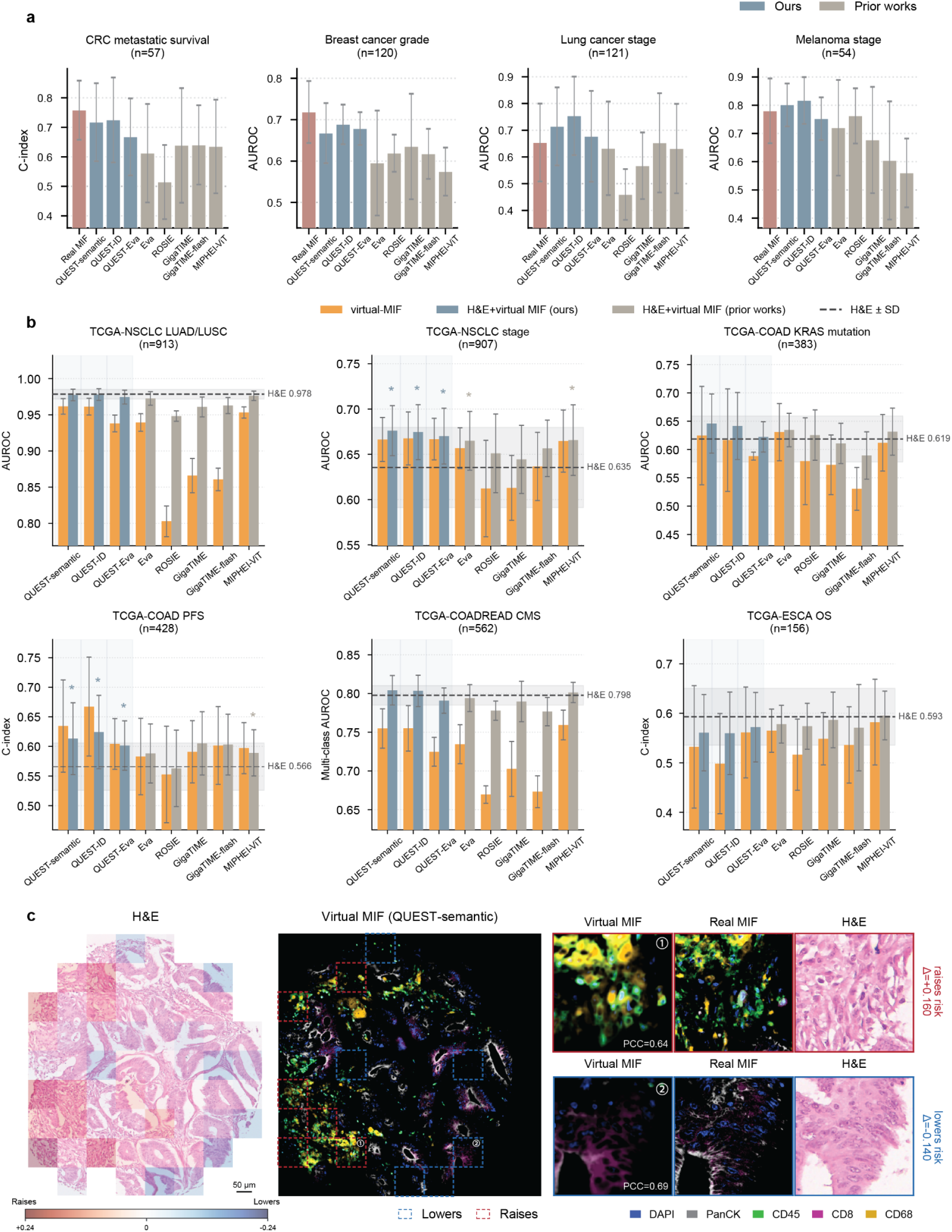
Downstream clinical utility of virtual MIF. **a**, Performance of real MIF and virtual MIF generated by our models or prior methods across 4 held-out paired H&E–MIF datasets, encompassing colorectal cancer metastatic survival, breast cancer grade, lung cancer stage, and melanoma stage. Survival performance was evaluated using the concordance index (C-index), whereas classification performance was evaluated using the area under the receiver operating characteristic curve (AUROC). **b**, Performance across 6 TCGA tasks using virtual MIF alone or virtual MIF combined with H&E. Dashed lines and associated shaded bands indicate the mean ± s.d. performance of the H&E-only baseline. Asterisks indicate significant improvements over the H&E-only baseline with bootstrap (**Methods**). In both panels, bars and error bars represent the mean ± s.d. across fivefold cross-validation. **c**, Patch-level contributions to a virtual MIF-based survival risk prediction in the metastatic CRC cohort. The contribution map highlights regions that increase or decrease predicted risk in red and blue, respectively. The corresponding whole-region virtual MIF is shown alongside zoomed-in examples of representative risk-raising and risk-lowering patches, with dashed boxes marking the top 10% of each. Zoomed-in views compare the virtual MIF, real MIF, and H&E images of risk-raising and risk-lowering patches.

We further evaluated the value of virtual MIF in complementing H&E-based predictions using public TCGA cohorts. The benefit of combining virtual MIF with H&E varied across tasks. For TCGA-NSCLC stage classification, H&E enhanced with QUEST-generated virtual MIF significantly improved upon the H&E-only baseline. Similarly, QUEST models yielded significant improvements in TCGA-COAD’s progression-free survival prediction over the H&E-only baseline. In TCGA-NSCLC Lung Adenocarcinoma/Lung Squamous Cell Carcinoma (LUAD/LUSC) classification, H&E alone achieved an AUC of 0.978, and only marginal gains were observed from virtual MIF. Modest gains were observed for TCGA-COAD KRAS mutation prediction and consensus molecular subtype (CMS) classification, whereas performance for TCGA-ESCA overall survival remained similar to the H&E-only baseline (**Figure 4b**). These results show that virtual MIF can improve H&E-based prediction for selected clinical endpoints, although the magnitude of benefit remains highly task-dependent.

To identify the driving factors of survival predictions, we quantified patch-level contributions by ablating each patch and measuring the resulting change in predicted risk. In the held-out metastatic CRC survival cohort, risk-raising regions showed stromal morphology accompanied by prominent CD68-associated signals in both virtual and real MIF, whereas risk-lowering regions were more tumor-rich (**Figure 4c** and **Supplementary Figure 12a**). In TCGA-COAD progression-free survival, risk-increasing patches appeared predominantly tumor-rich, whereas risk-decreasing patches showed greater immune infiltration (**Supplementary Figure 12b-c**, Williams et al. 2024). These distinct attribution patterns may reflect differences in clinical state and outcome definition. In the metastatic cohort, where all patients had established metastatic disease, stromal, CD68-rich risk regions may reflect a macrophage-associated immunosuppressive microenvironment, consistent with the adverse prognostic relevance of stromal programs and specific tumor-associated macrophage states in CRC (Joanito et al. 2022; Yedan Liu et al. 2022). In the more clinically heterogeneous TCGA-COAD cohort, contrasts between tumor-dominant and immune-infiltrated regions may remain the major informative correlate of progression risk.

Motivated by the tumor–immune contrast observed in the patch-level contribution maps, we next asked whether individual virtual marker signals were associated with patient outcomes in TCGA-COAD. Kaplan–Meier analyses showed that higher virtual CD68, CD8, and CD20 signals were associated with longer progression-free survival, whereas virtual DAPI and PanCK were not associated with prognosis (**Supplementary Figure 13a**). The favorable association observed for CD8, together with the broader prognostic relevance of quantitatively assessed immune infiltration, is consistent with large-scale studies of CRC trial cohorts (Frei et al. 2024). We further evaluated the virtual marker signal CD8/PanCK ratio as a simple, interpretable, and MIL-independent feature representing cytotoxic T-cell infiltration relative to epithelial tumor content. A higher CD8/PanCK ratio significantly stratified progression-free survival in Kaplan–Meier analysis (log-rank P = 0.010) and remained independently protective in multivariable Cox regression after adjustment for the virtual-MIF MIL score, age, sex, stage, and MSI status. The MIL score also remained independently associated with increased risk, indicating that it captured prognostic information complementary to the CD8/PanCK ratio (**Supplementary Figure 13b**).

### Cross-modal retrieval bridged by virtual MIF staining

In clinical practice, MIF is rarely acquired, which limits its direct applications in clinical decision-making for new patients. However, the growing number of public and institutional collections makes it increasingly feasible to assemble large-scale MIF reference galleries of tissues (HuBMAP Consortium 2019; Rozenblatt-Rosen et al. 2020). Such galleries could potentially provide molecular contexts for H&E-stained samples by linking them with existing MIF references when their exact counterparts are unavailable. We therefore investigated whether virtual MIF staining could serve as the bridge between the two modalities by mapping an H&E query into MIF space and retrieving reference MIF patches with similar protein expression patterns and cellular composition.

First, we used paired H&E–MIF patches from the held-out Mixed 10K dataset. For each H&E query, the patch was translated into virtual MIF, which was then encoded using a frozen foundation model (Yufan Liu et al. 2025) to obtain the query embedding. Candidate patches from the MIF gallery were encoded through the same procedure and ranked by their cosine similarity to the query (**Figure 5a)**. The gallery comprised the MIF counterparts of all eligible patches in Mixed 10K, with each paired patch serving once as an H&E query during evaluation. Patches from the query’s source region were excluded from the gallery. We evaluated the retrieval efficacy of different virtual MIF staining models along with a specialized cross-modal retrieval model, Haiku (Cui et al. 2026), using similarity metrics in the protein expression and cellular composition space. QUEST models produced concordant retrievals among the evaluated approaches. QUEST-semantic and QUEST-ID achieved a top-1 expression cosine similarity of 0.52, compared with 0.15–0.37 for previous virtual MIF staining methods and 0.38 for Haiku (**Figure 5b**). In addition, QUEST-semantic yielded the lowest L2-distance distribution, which differed significantly (p < 0.001) from those of all other methods except QUEST-ID by the Kolmogorov–Smirnov test (**Supplementary Figure 14a**). Consistently, QUEST models achieved the highest precision profiles across retrieval depths, with QUEST-semantic generally performing the best (**Supplementary Figure 14b**).

**Figure 5:**
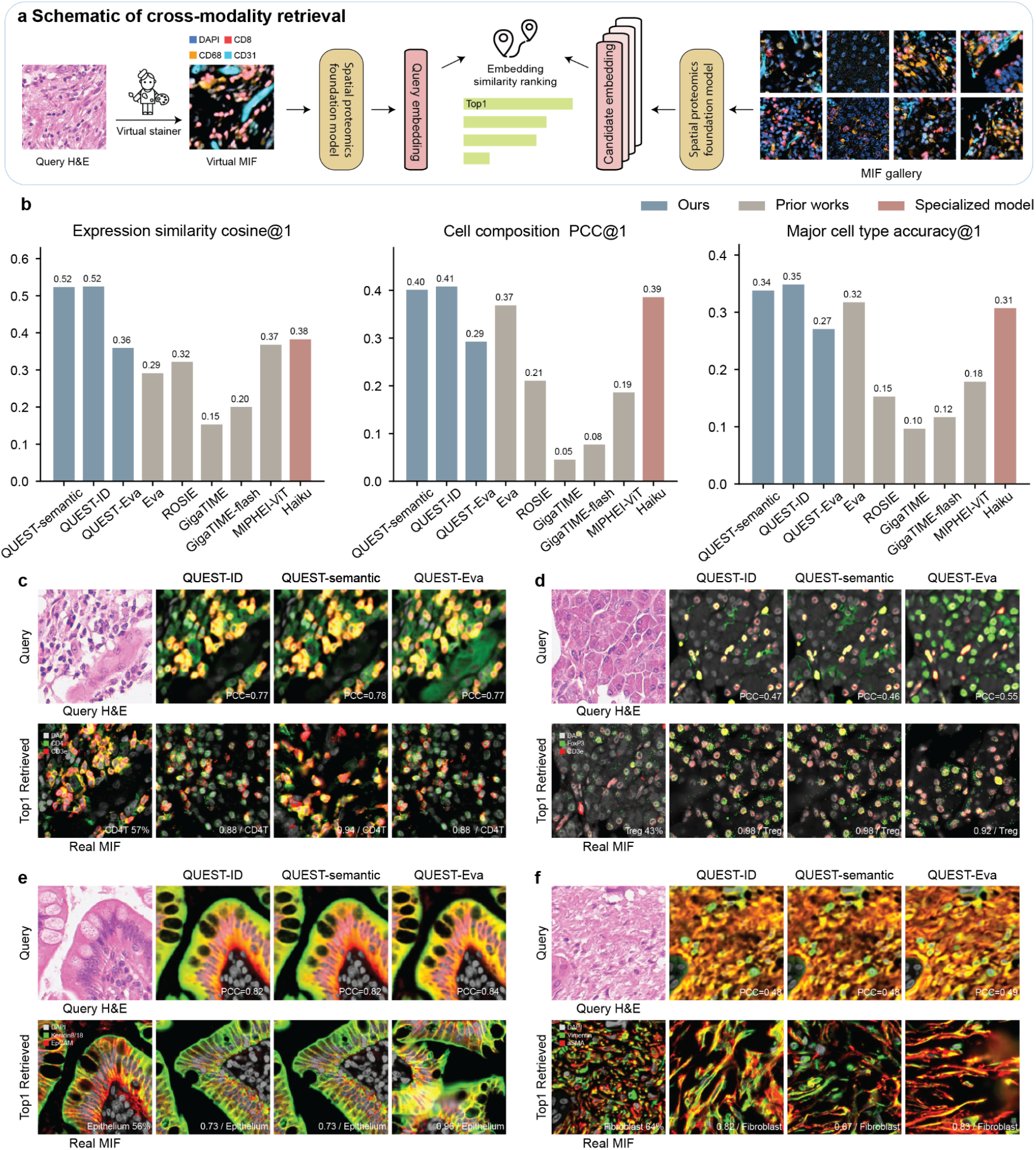
Cross-modality image retrieval through virtual staining. **a**, Schematic of the retrieval workflow. A virtual staining model predicts MIF from a query H&E image, and a spatial proteomics foundation model, Eva, encodes the virtual MIF into a query embedding. Real MIF images in the reference gallery are encoded using the same foundation model, and candidates are ranked by embedding similarity. **b**, Top-1 retrieval performance evaluated by the cosine similarity of protein expression profiles, PCC between cell compositions, and accuracy of the dominant cell type. The real MIF paired with each query H&E image was used only for evaluation and did not participate in candidate ranking. **c–f**, Retrieval examples dominated by CD4 T cells (**c**), regulatory T cells (**d**), epithelial cells (**e**) and fibroblasts (**f**). For each example, the top row shows the query H&E image and virtual MIF predictions from QUEST-ID, QUEST-semantic, and QUEST-Eva, and the bottom row shows the paired real MIF reference and the top-1 real MIF image retrieved by each model. Values overlaid on virtual MIF images indicate PCC with the paired real MIF. Percentages on the paired real MIF indicate the abundance of the dominant cell type, whereas values and labels on retrieved images indicate cell composition PCC and the dominant cell type, respectively.

We next examined whether the retrieved patches also recapitulated the cellular compositions. This analysis used regions from the held-out test data, comprising gastrointestinal stromal tumor and pancreas cancer tissues with cell segmentation and cell-type annotations derived from MIF images (**Methods**). Similarly, QUEST-semantic and QUEST-ID achieved top-1 cell-composition PCCs of 0.40–0.41, exceeding Haiku (0.39), Eva (0.37), and the other previous methods (0.05–0.21) (**Figure 5b**). QUEST-semantic and QUEST-ID also obtained the highest top-1 major cell type accuracy of 0.34–0.35, respectively, exceeding Haiku (0.31), Eva (0.32), and the other previous methods (0.10–0.18). Visual inspection of the top-ranked candidates was consistent with the quantitative results. For the displayed H&E query, QUEST models prioritized MIF patches that closely matched the query’s measured marker expression profile and cell type composition and also shared its dominant cell type and tissue category (**Figure 5c-f, Supplementary Figure 14c-e**).

## Discussion

Inferring spatial protein expression from routine H&E could extend molecularly informed tissue analysis to cohorts where direct measurements of molecular data are impractical. Existing virtual MIF staining methods are typically restricted to predefined marker panels, and their utility for downstream biological and clinical analyses remains insufficiently understood. In this study, we developed QUEST, a class of models that dynamically take the requested markers as query inputs and generate their corresponding expression intensity maps. This query-based formulation allows a single model to accommodate heterogeneous marker panels rather than being tied to a predefined output panel. We evaluated the resulting virtual MIF across biological scales, from pixels and cells to tissue regions and patients, as well as heterogeneous marker panels and imaging platforms.

Multi-scale evaluations suggested that virtual MIF staining cannot be adequately assessed by a single image-level metric. In particular, our analysis exposed a failure mode of SSIM (Wang et al. 2004) in sparse MIF images, where empty predictions received deceptively high scores. We therefore complemented standard similarity and correlation measures with foreground intensities and foreground-background discrimination analyses. Across metrics, QUEST generated virtual MIF that achieved strong agreement with real MIF in held-out and external datasets, reconstructing signals across diverse tissue categories and marker panels. We further showed that pan-cancer training improved tissue-specific prediction and supported cross-tissue transfer of marker supervision. Virtual MIF preserved relative marker expression intensities and marker-positive populations at the cell level, supporting multi-marker cell typing, cellular neighborhood profiling, and characterization of their spatial organization, such as T-cell proximity to vascular structures and local cell-type diversity. Notably, the end-to-end workflow based on virtual DAPI and virtual marker expression reproduced major cellular neighborhood patterns. These findings demonstrate the biological fidelity of virtual MIF captured through a multi-scale evaluation of marker-positive signals, cellular composition, and tissue organization.

At the patient level, virtual MIF could retain clinically relevant signals captured by real MIF and could augment H&E-based prediction in the absence of molecular signals. Concretely, virtual MIF is inferred from H&E, and these improvements should not be interpreted as recovery of independent molecular measurements. Instead, virtual MIF staining exposes protein-associated morphological patterns to downstream predictions. Patch-level contribution maps further localized stromal, tumoral, and immune-infiltrated regions associated with predicted risk in the held-out metastatic colorectal cancer and TCGA-COAD survival cohort. Consistently, Kaplan-Meier analyses identified associations between virtual immune marker signals and progression-free survival, where the virtual CD8/PanCK ratio was independently associated with better prognosis. These analyses show that virtual MIF staining can provide spatially and biologically interpretable evidence for clinically relevant associations, beyond improving predictive performance alone.

We demonstrated cross-modal retrieval as an application of virtual MIF staining. By mapping an H&E query into the MIF representation space, QUEST enabled retrieval of MIF patches with similar protein expression profiles and cellular compositions. Notably, QUEST’s retrieval performance matched or exceeded the dedicated model, Haiku, without requiring task-specific alignment or training. In clinical settings, when matched spatial protein expression measurements are unavailable, this framework could provide molecular context for interpreting H&E-only inputs and help prioritize subsequent analysis.

QUEST models can generate different combinations and numbers of supported markers at inference without panel-specific retraining. In addition, QUEST maintained reasonable GPU memory use and inference time when scaling to hundreds of markers (**Supplementary Figure 15**). This provides a practical path toward larger marker panels and may support the extension of query-based prediction to high-dimensional spatial multi-omics maps.

The three QUEST variants differed in their visual pretraining and marker representations. QUEST-ID and QUEST-semantic use a pathology foundation model to encode H&E and represent the requested markers using learned identity embeddings or fixed text-derived embeddings, respectively. QUEST-Eva instead adapts a model pretrained jointly on spatial proteomics and matched histopathology, with marker embeddings conditioning the reconstruction of requested channels. Whereas QUEST-ID is restricted to markers encountered during training, QUEST-Eva and QUEST-semantic can construct queries from externally derived marker embeddings, providing a potential route beyond the training marker vocabulary.

To evaluate marker-level extensibility, we excluded 24 markers entirely from QUEST-semantic training and tested whether the semantic query formulation could generalize beyond the training marker vocabulary. In this zero-shot setting, test markers were excluded entirely from training and introduced only through their semantic descriptions at inference. Zero-shot QUEST-semantic retained 56–60% of the correlation achieved under full supervision, demonstrating partial transfer to previously unseen markers (**Supplementary Figure 16**). Notably, the predictive performance was also sensitive to marker prompting (**Methods**), with function- and drawing-based descriptions emphasizing different aspects of marker identity, indicating the need to carefully formulate the biological information used to represent markers.

We further applied QUEST to a prostate cancer sample from a nondestructive 3D pathology dataset (Xie et al. 2022) as a qualitative proof-of-concept. Applied across successive optical planes of the volumetric H&E image, QUEST-semantic generated virtual MIF predictions that could be assembled into a coherent 3D representation (**Supplementary Figure 17, Supplementary Video 1**). The reconstructed CD31 signal clearly delineated vascular structures and their continuity across tissue depth. Such volumetric virtual staining could support more complete cell phenotyping and enable the analysis of tissue architecture, cell–cell contacts, and multicellular immune niches throughout intact tissue volumes (Yapp et al. 2025) in future research.

Several limitations remain. Although the training atlas consisted primarily of PhenoCycler and PhenoCycler Fusion data, external evaluations on ORION-CRC, HEMIT, and HuBMAP-intestine provided preliminary evidence of generalization across additional imaging platforms and staining protocols. These external datasets nevertheless remain limited in scale and marker coverage, and larger cross-platform evaluations will be needed to further establish the robustness of virtual MIF. Evaluations of virtual MIF prediction are bounded by the quality of experimental ground truth, and future benchmarking and model development will benefit from curated, gold-standard datasets with stringent quality control and autofluorescence/noise correction to establish the true limits of virtual MIF staining fidelity. Prediction also remains bounded by the molecular information reflected in H&E. Our per-marker evaluations and independently trained single-marker U-Net references showed substantial variation in predictability across proteins, with markers associated with visually distinct cellular or tissue structures generally being more predictable than sparse immune or functional markers. Variations in virtual MIF prediction performance were observed across different tissue types and were not fully explained by per-tissue training data volume, further indicating heterogeneous, tissue-dependent influences from morphology, marker composition, and intrinsic marker predictability.

To surpass these bottlenecks while maintaining clinical feasibility, integrating accessible modalities such as low-plex immunohistochemistry or bulk transcriptomics during prediction may be a promising path towards revealing the nuanced molecular states. Further, future work may better assess and improve generalization across MIF platforms and provide confidence estimates for individual marker predictions. These advances will help determine which virtual markers are sufficiently reliable for downstream analysis and which still require experimental validation.

The patient-level analyses presented here should be viewed as proof-of-concept evaluations of clinical utility, and further validation in larger, independent external cohorts with paired H&E–MIF assays and well-annotated outcomes will be required. Moreover, while we demonstrated that virtual MIF can support cell typing and CN annotation, deploying it for larger-scale discovery of novel disease-associated tumor-microenvironment biomarkers remains a challenging and resource-intensive application reserved for future investigations.

Overall, we established QUEST to transform virtual MIF staining from fixed-panel image translation into a flexible interface between H&E-based histology and spatial proteomics. QUEST dynamically generated virtual MIF intensity maps for the queried markers, which preserved information relevant to cellular phenotyping, tissue organization, patient-level analysis, and cross-modal retrieval. With broader prospective validation and reliable uncertainty estimation, QUEST could transform vast H&E archives into queryable, molecularly informed resources, enabling population-scale mapping of tissue ecosystems, accelerating biomarker discovery, and extending spatially informed precision medicine far beyond the scale currently accessible to direct MIF profiling.

## Methods

### Datasets

#### Atlas for model development

In this study, we curated a large-scale paired H&E–multiplex immunofluorescence (MIF) atlas across marker panels and tissues. The atlas contains 12,880 tissue regions with resolution of 0.3775–0.5 µm/pixel from 21 tissue categories and collectively covers 129 protein markers. Most regions were generated using the Akoya PhenoCycler platform. Each region has a matched H&E image and a region-specific MIF panel. Marker panels are not identical across samples, with a median of 53 measured markers per region (**Supplementary Figure 1a-b**). Model inputs and targets were represented as spatially paired non-overlapping 224×224 patches, with pixel intensities normalized to [0, 1] independently for each marker within each tissue region. The split was performed at the patient level within each tissue category. A total of 11,068 regions were assigned to model development, comprising 9,791 training regions and 1,277 validation regions. The remaining 1,812 regions formed the held-out evaluation pool. The indexed training, validation, and test portions contained 1,926,041, 213,943, and 260,352 paired patches, respectively.

#### Image-level evaluation datasets

All held-out image fidelity evaluation datasets were derived from the held-out test regions described above. None of these H&E images, MIF images, or clinical labels were used to optimize models in this study. The multi-tissue Mixed 10K benchmark was formed by sampling 10,000 patches from the standard atlas test split and contained 52 evaluable markers. For the tissue-specific benchmarks, a fixed set of 2,000 patches was sampled from each pool to form the Breast-stain, CRC-stain, melanoma-stain, and Lung-stain datasets, with 54, 57, 67, and 78 markers evaluable, respectively. The same sampled patches and measured targets were used for every virtual staining model. For region- and tissue-level evaluations, up to 100 held-out regions were sampled per tissue type. Images in these analyses had spatial resolutions ranging from 0.3775 to 0.5 µm/pixel.

External image-level evaluations used paired H&E and multiplex-staining datasets that were not part of the atlas for model development or held-out datasets. HEMIT (Bian et al. 2024) is a public paired H&E and multiplex-immunohistochemistry benchmark generated using the PANO 7-plex mIHC workflow. The evaluation contains 8 whole-slide images with a resolution of around 0.5 μm/pixel, stained with three markers: DAPI, PanCK, and CD3e. ORION-CRC (J.-R. Lin et al. 2023) is a public colorectal cancer cohort acquired on the RareCyte ORION platform, and the evaluated subset contains 41 whole-slide images with a resolution of 0.5 μm/pixel, covering 17 biomarkers. ORION-CRC was used to train MIPHEI-ViT, so MIPHEI results on this cohort are identified as in-domain results. HuBMAP-intestine (Hickey et al. 2023; HuBMAP Consortium 2019) is a public intestinal dataset with a resolution of 0.3775 μm/pixel, acquired using the first-generation CO-Detection by indEXing (CODEX) platform with a Keyence BZ-X700 microscope, followed by H&E staining of the same tissue section. After registration and quality-control filtering, we retained 4 slides with 44 markers stained from a single donor for evaluation.

#### Cell-level evaluation datasets

The atlas-derived cell-level benchmark comprised 114 regions from the held-out pool for which cell segmentation masks were available. These regions contained 14,134 patches and 1,159,047 unique segmented cells. The external cell-level dataset was the Schürch (Schürch et al. 2020) colorectal cancer CODEX cohort, accessed through the PathoCell (Lüscher et al. 2026) representation with a resolution of 0.3775 μm/pixel, which provides matched H&E, cell segmentation masks, and cell type labels. The full analysis used 109 tissue regions and 220,977 unique segmented cells. For cellular neighborhood (CN) analyses, we used 16 held-out esophagus regions containing 135,281 segmented cells and 20 held-out lung regions containing 455,231 segmented cells.

#### Clinical cohorts for downstream tasks

Clinical cohorts were assembled by joining the held-out region index to the corresponding patient annotations. The CRC (colorectal cancer) metastatic survival cohort with resolution of 0.5 µm/pixel contained 62 regions from 57 unique patients, corresponding to 37 patient deaths. The breast cancer grade cohort with resolution of 0.5 µm/pixel included 124 regions from 120 patients, with 34 high-grade and 86 low-grade patients. The lung cancer stage cohort included 121 patients, with 25 stage III–IV and 96 stage I–II cases, with resolution of 0.3775–0.5 µm/pixel. The melanoma stage classification used 56 regions from 54 patients, with 29 patients of stage III–IV disease and 25 of stage I–II disease, with resolution of 0.3775–0.5 µm/pixel.

External downstream tasks were evaluated on The Cancer Genome Atlas (TCGA, Cancer Genome Atlas Research Network et al. 2013) datasets with a resolution of around 0.25–0.5 μm/pixel. TCGA slides do not have paired MIF in this study, and virtual MIF was generated directly from each diagnostic H&E slide. The TCGA-COAD (Cancer Genome Atlas Network 2012) cohort contained 428 patients for progression-free survival. Each survival endpoint comprised a follow-up time and an event indicator, with death defining the overall survival event and progression or death defining the progression-free event, and patients without an event were censored. The classification analyses included 383 patients for KRAS status, classified as KRAS mutant or KRAS wild type. The TCGA-COADREAD (Joanito et al. 2022) cohort combined TCGA-COAD and TCGA-READ and was used for 4-class consensus molecular subtype prediction. The final analysis included 562 patients, comprising 418 from TCGA-COAD and 144 from TCGA-READ. The TCGA-NSCLC (Cancer Genome Atlas Research Network 2012) cohort combined TCGA-LUAD and TCGA-LUSC. Histological subtypes were evaluated in 913 patients, with the histology labels being lung adenocarcinoma (LUAD) and lung squamous-cell carcinoma (LUSC). Pathological stage was available for 907 patients and was binarized as stage I versus stages II–IV. The TCGA-ESCA (Cancer Genome Atlas Research Network et al. 2017) cohort contained 156 patients, and the overall survival endpoint distinguished observed deaths from censored follow-up while retaining the survival time.

### Model development and training

#### Model architecture of QUEST

We proposed QUEST (**QUE**ry-based virtual **STaining)**, a parameter-efficient deep learning model for predicting multiplex immunofluorescence (MIF) images from hematoxylin and eosin (H&E) images using dynamic text-based marker embedding queries. For an RGB H&E patch *x*^*H&E*^ ∈ [0, 1] ^3×224×224^ and a set of requested markers *S* = (*m*_1_,…, *m* _*K*_), the model produces *Y* ^*MIF*^= *f* _θ_ (*x* ^*H&E*^, *S*) ∈ ℝ ^*K*×224×224^. Here *K* is the requested marker panel size. The model consists of a frozen H&E encoder, a shared marker query projector, a multi-scale H&E decoder, a cross-attention marker-to-H&E readout module, and a predictive output head (**Supplementary Figure 2a**).

##### H&E encoding

H&E features were obtained using the pretrained histopathology foundation model UNI2-h (R. J. Chen et al. 2024) encoder. For a 224×224 input H&E patch normalized to [0,1], each 14×14 pixel token was encoded in a 1,536-dimension feature, forming a feature as *Z*_*H&E*_ = *E*_*UNI*2−*h*_(*x*^*H&E*^) ∈ ℝ^256×1536^. The resulting spatial token representations were used in two parallel branches, namely the marker-to-H&E cross-attention readout module and the multi-scale H&E decoder.

#### Shared marker query projector

Each marker m was represented by an initial vector 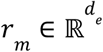. QUEST-ID and QUEST-semantic differed in how their initial marker representations were constructed. QUEST-ID used a learnable marker identity lookup table, which assigned an independent 4,096-dimensional vector to each marker in the training vocabulary. By contrast, QUEST-semantic represented each marker using a frozen 3,072-dimensional embedding of its textual description generated by OpenAI’s text-embedding-3-large model (https://developers.openai.com/api/docs/models/text-embedding-3-large) and did not use a learnable marker identity table. The textual description of markers is listed in **Supplementary Data 1**. The marker embeddings in QUEST-semantic were normalized and mean-centered over all markers in training. In both variants, the resulting marker query representation was mapped into the 512-dimensional decoder space using a projection shared across markers, which was denoted as *M* ∈ ℝ ^*K*×512^. As each marker is processed independently through shared model parameters, both QUEST-semantic and QUEST-ID support arbitrary combinations of markers from their respective supported vocabularies. Notably, QUEST-semantic additionally supports zero-shot prediction of previously unseen markers.

#### Marker-to-H&E cross-attention

Marker embedding and H&E embedding are fused in a query-based approach. The 1,536-dimensional H&E feature was first projected to the same dimension as the marker queries, denoted as *H* ∈ℝ ^256×512^. We used an 8-layer cross-attention (Vaswani et al. 2017) decoder in which the requested marker embeddings served as queries and the spatial H&E tokens served as keys and values. For the layer *l* and attention head *h* ∈ {1,…, 8}, the attention map was computed, and the H&E-informed marker features were computed as

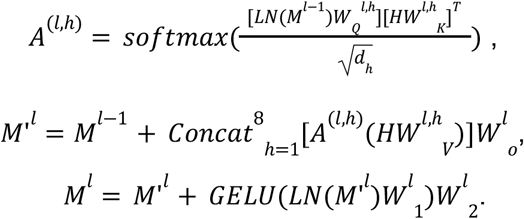

Here,*M*^*l*−1^ ∈ ℝ^*K*×512^ denotes the *K* marker queries entering the layer *l* and *H* ∈ ℝ ^256×512^ denotes the projected H&E token representation, which are described above.*LN* is the LayerNorm operation (Ba et al. 2016). We used 8 heads with a dimension *d* _*h*_ of 64, with *W* _*Q*_, *W* _*k*_, *W*_*V*_ ∈ ℝ^512×64^, and *W*_*O*_ ∈ ℝ^512×512^ as an output projection. The feed-forward network (FFN) expanded the feature dimension from 512 to 2,048 before projecting it back to 512 dimensions. Each decoder layer used a pre-normalized residual design, following residual learning (He et al. 2015). GELU (Hendrycks and Gimpel 2016) was used as the activation function. Each marker query was updated independently using the shared spatial H&E representation without marker-marker interaction.

##### Multi-scale H&E decoder

To recover fine-grained spatial information, we implemented a coarse-to-fine, multi-scale decoder with lateral feature fusion (T.-Y. Lin et al. 2016). The spatial token representation *Z* _*H&E*_ from UNI2-h was first reshaped to a feature map of size 1,536×16×16 and projected to a channel number of 256. In parallel, the original H&E image was processed by a frozen ConvNeXt-Base (Z. Liu et al. 2022) backbone to extract a hierarchy of convolutional feature maps. The four ConvNeXt stages produced feature maps with spatial resolutions of 56×56, 28×28, 14×14, and 7×7, containing 128, 256, 512, and 1,024 channels, respectively. Each feature map was independently projected to 256 channels using a 1×1 convolution. Whereas the UNI2-h tokens provided high-level histopathological context, the ConvNeXt feature hierarchy retained progressively finer local information about cellular morphology and tissue boundaries.

The projected UNI2-h feature map was resized to 7×7 and fused with the coarsest ConvNeXt feature map through channel concatenation and convolution. The fused representation was then progressively upsampled and combined with lateral ConvNeXt features at 14×14, 28×28, and 56×56. The final representation was resized and convolutionally refined to produce a dense H&E feature map *F*_*H&E*_ ∈ ℝ^256×224×224^. The ConvNeXt backbone remained frozen, whereas the multi-scale decoder was trained jointly with QUEST.

##### Predictive output head

We denoted the final H&E-informed marker query features produced by cross-attention as *F*_*marker*_ ∈ ℝ ^*K*×512^ and K is the number of requested markers.

To generate the final marker-specific intensity maps of prediction, we applied feature-wise linear modulation (Perez et al. 2018), and the feature of each marker was first projected into a readout vector *r*_*k*_ and modulation parameters γ_*k*_ and β_*k*_, each with 256 dimensions. The intensity at a spatial position *h, w* is then predicted as:

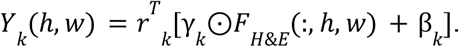

Here ⊙ denotes element-wise multiplication. Thus, each marker feature selectively modulated the shared H&E representation to generate its corresponding continuous MIF intensity map. The projection layers were shared across markers, allowing arbitrary marker sets to be requested at inference.

### Fine-tuning of QUEST-Eva

QUEST-Eva was obtained by fine-tuning the pretrained Eva (Yufan Liu et al. 2025) masked autoencoder following the H&E-to-MIF adaptation procedure described in the Eva study. To ensure consistency with the data partition used in this study and prevent information leakage, we repeated the fine-tuning using only our designated training set, with checkpoint selection performed on the corresponding validation set. All model inputs, channel embeddings, masking strategy, reconstruction objective, and inference procedure were otherwise identical to those described in Eva. Briefly, the RGB H&E images were treated as separate channels and concatenated with multi-channel MIF images, with channel identities encoded using the pretrained GenePT (Y. Chen and Zou 2025) marker embeddings. All MIF spatial tokens were masked while the 3 H&E channels remained visible, and reconstruction loss was computed only over the masked MIF targets. The pretrained initialization checkpoint is available at https://huggingface.co/yandrewl/Eva.

### Model training and fine-tuning

In the training of QUEST, UNI2-h was frozen during training, and the H&E features were cached in practice so that the H&E encoder was not instantiated. For each patch, a maximum number of 30 markers were randomly sampled from the channels measured in that region and padded for batching. Consequently, the marker panel and channel ordering varied among training examples. QUEST was optimized with AdamW (Loshchilov and Hutter 2017), a 300-step linear warmup, and cosine learning rate decay from 3×10^-4^ to 10^-6^. QUEST-ID and QUEST-semantic were both trained with a batch size of 160. Training converged after approximately 50,000 steps. We trained QUEST using a signal-weighted L1 loss, specifically 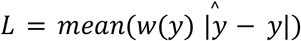 and 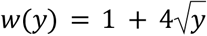. This weighting was designed to prevent the large proportion of background pixels from dominating optimization while retaining supervision over background regions.

QUEST-Eva was initialized from the publicly released pretrained Eva model and fine-tuned on the same paired H&E–MIF collection using the same data splitting as QUEST. Following the H&E-to-MIF adaptation procedure introduced for Eva, QUEST-Eva was fine-tuned for 20 epochs by masking the MIF channels and reconstructing them from the visible H&E channels using mean squared error (MSE) as reconstruction loss.

The numbers of trainable parameters, excluding frozen components, were 39.75 million for QUEST-semantic, 41.08 million for QUEST-ID, and 122.44 million for QUEST-Eva.

### Evaluation tasks and metrics

All model selection and hyperparameter tuning were performed using the training and validation partitions. The final weights were evaluated on held-out data or external cohorts that were not used for model optimization. Marker names were harmonized to the QUEST vocabulary before comparison, with only straightforward nomenclature aliases resolved, such as Hoechst to DAPI, CD8a to CD8, Pan-CK/CK to PanCK, SMA to aSMA, E-cadherin to ECad, PD-1 to PD1, PD-L1 to PDL1, and FOXP3 to FoxP3. No biological proxy substitutions were made. Unless otherwise stated, image-level metrics were calculated separately for each marker within each patch, averaged across markers to obtain a patch-level score, and then averaged across patches to obtain the dataset-level result.

#### Pixel-, region-, and tissue-level virtual staining fidelity

We first quantified agreement between each continuous virtual MIF prediction and its real MIF reference at various scales. At the patch level, Pearson’s correlation coefficient (PCC) and Spearman’s rank correlation coefficient (Spearman’s ρ) were calculated between predicted and real pixel intensities for each marker. To assess whether the models recovered marker expression within local tissue regions, we additionally performed a grid-level evaluation. Each predicted and measured marker image was partitioned into a 14×14 spatial grid, and pixel intensities were averaged within each grid cell. PCC and Spearman’s ρ were then calculated between the corresponding predicted and measured grid-cell values for each marker. For the per-marker analysis on Mixed 10K, PCC values were instead averaged across patches separately for each marker and reported by biological group.

To estimate a marker-specific reference for the signal recoverable from H&E morphology, we trained an independent supervised U-Net (Ronneberger et al. 2015) for each of the 20 markers spanning major cellular compartments and biological functions. For this analysis, regions from the original QUEST test partition were divided into disjoint U-Net training and evaluation subsets. Each U-Net was trained only on the U-Net training regions, and both the U-Net and QUEST were evaluated on the same patches from the evaluation regions, which had been held out from the training of both models. U-Nets were trained for 12,000 steps using AdamW with a learning rate of 2×10^-4^. As these marker-specific U-Nets received direct supervision from other regions in the test split, whereas our models had never been trained on any region from that split, their performance was interpreted as a predictability reference.

Structural similarity was assessed with the structural similarity index measure (SSIM, Wang et al. 2004). As sparse MIF images contain large background areas, we also evaluated an all-zero image as a whole-black baseline. This control quantifies the SSIM that can be obtained without recovering any marker-positive signal. SSIM was therefore treated as a complementary structural metric rather than as the sole measure of virtual staining fidelity.

To address this, we performed two evaluations of marker-positive regions. First, an Otsu (Otsu 1979) threshold was applied to the real MIF markers to define reference foreground pixels. Foreground PCC was then calculated between real and predicted intensities using only positive pixels based on real MIF images. Second, foreground-background discrimination was treated as a binary prediction task. The Otsu-derived measured-MIF mask served as the ground truth. For F1-score calculation, each continuous prediction was binarized by Otsu thresholding, and for area under the precision-recall curve (AUPRC), the predicted intensities were used as scores against the binary reference labels. These analyses jointly measure recovery of positive signals, missed foregrounds, and spurious signals in the background.

As the evaluated models supported different marker panels, we additionally performed shared marker comparisons. For each prior method, these comparisons were restricted to the intersection of markers supported by that method and by our models. The number of common markers included in each comparison is reported in the corresponding figures.

To assess generalization beyond individual patches, we further evaluated up to 100 held-out tissue regions per category across 21 tissue categories. PCC was calculated between the predicted and real signal for each available region-marker pair, and scores were averaged within each tissue category. A model-level tissue score was obtained by averaging across tissue categories. Whole-slide image predictions on held-out and TCGA slides were used as qualitative demonstrations of spatial scalability.

#### Pan-cancer training and cross-tissue marker transfer

To assess the effect of multi-tissue training, we compared QUEST-semantic models trained across all tissue categories with single-tissue models trained only on colon/rectum or lung. The full-panel comparison included 57 markers for colon/rectum and 78 markers for lung. To evaluate cross-tissue marker transfer, PDL1, FOXP3, Ki67, BCL2, and EpCAM were removed from all training and validation regions of the target tissue. Under this marker-holdout setting, the multi-tissue model retained supervision for these markers from other tissue categories, whereas the single-tissue model received no direct supervision for them.

Within each comparison, models were trained from scratch using identical architectures and optimization settings to QUEST-semantic. Performance was evaluated using per-marker full-image PCC and averaged across either the complete target-tissue panel or the five held-out markers. Paired differences were assessed using a paired region-level bootstrap, in which test regions were resampled rather than individual patches.

#### Cell-level evaluation

Cells were segmented with Mesmer (Greenwald et al. 2022) from the DAPI channels of real MIF. The resulting cell masks were applied identically to the real and predicted MIF images. For each cell and marker, the continuous cell-level signal was defined as the mean pixel intensity within the corresponding cell mask. For the continuous cell-level fidelity analysis, PCC between real and predicted cell-level intensities restricted to the cell mask was calculated for each region–marker pair and then averaged across regions to obtain the per-marker result.

We additionally evaluated recovery of marker-positive cells as a binary task. A marker- and region-specific percentile-normalized threshold was applied to the measured cell-level intensities to assign each cell a reference positive or negative label, whereas the continuous virtual MIF cell intensities were retained as prediction scores. Cell-level discrimination was quantified by AUROC, which evaluates the ranking of positive versus negative cells across all possible decision thresholds. For visualization, thresholding the real signal yielded K positive cells for a given marker and region. The real MIF map displays these K positive cells, while each virtual MIF map displays the K cells with the highest predicted intensities.

We evaluated cell segmentation from virtual DAPI and H&E-derived hematoxylin. Real and virtual DAPI images were independently segmented using the same Mesmer configuration. For H&E segmentation, RGB images were converted to optical-density space and separated by color deconvolution using a fixed hematoxylin–eosin stain matrix. The extracted hematoxylin channel was then processed using the same Mesmer configuration. Virtual DAPI- and H&E-derived masks were evaluated against the real DAPI-derived masks. Predicted–reference instance pairs were selected in descending order of IoU, with each instance matched at most once. Pairs with IoU ≥ 0.5 were counted as true positives, whereas unmatched predicted and reference instances were counted as false positives and false negatives, respectively. Segmentation performance was reported using the F1 score at an IoU matching threshold of 0.5 (F1@IoU 0.5).

#### Cell typing and neighborhood analysis

We evaluated whether multichannel virtual MIF profiles supported downstream cell typing using the PathoCell (Lüscher et al. 2026) cohort. All models were evaluated on the same cells, segmentation masks, reference annotations, and region-disjoint data partition. As the models predicted different marker panels, each model was evaluated using the largest subset of measured PathoCell markers that it supported. A linear multinomial classifier was fitted to the resulting per-cell marker intensity vectors using the same cell-typing procedure and data partitions for real and virtual MIF.

Two related classification tasks were evaluated. The first used the native 15-class PathoCell annotation scheme. The second mapped the fine-grained annotations from the original cell annotation (Schürch et al. 2020) into 6 biologically broader classes: B lineage, CD4 T cells, CD8 T cells, epithelial cells, myeloid cells, and stromal/vascular cells. Performance for each taxonomy was summarized by the macro-F1 score and cell-level accuracy in visualization examples.

We next tested whether cell types inferred from virtual MIF preserved spatial relationships. For T cell to vascular proximity analysis, cell centroid coordinates were used to calculate the Euclidean distance from each T cell to its nearest cell classified as vasculature/lymphatics in 15-class annotations. This calculation was performed separately using the real MIF reference labels and the cell types inferred from each virtual MIF prediction. The resulting cumulative distance distributions and their median distances were compared.

Local cell-type diversity was quantified for every cell by calculating the Shannon entropy, in bits, of the cell-type distribution among its ten nearest neighboring cells. Ground truth entropy was calculated from real MIF-derived cell types, whereas predicted entropy was calculated from the corresponding virtual MIF-derived cell types using the same cells and neighbor graph. Agreement between predicted and ground-truth entropy was quantified by Spearman’s ρ across cells.

#### Cellular neighborhood annotation

Cellular neighborhood (CN) annotation followed a sequential workflow from MIF signals to cell types and subsequently from local cell type compositions to CN assignments. For the real MIF reference, cells were segmented from measured DAPI using Mesmer (Greenwald et al. 2022). Marker expression was calculated as the mean intensity within each cell mask and normalized to [0,1]. Cell types were defined by k-means clustering of the normalized real MIF profiles. For each marker, cluster centroids were converted to z-scores across cell types, and markers with z≥1 were considered enriched. Cell types were annotated according to their enriched marker combinations, with the resulting definitions provided in **Supplementary Table 1**. For each cell, a local composition vector was constructed from the frequencies of cell types among its 10 nearest spatial neighbors. These composition vectors were subsequently clustered by k-means to define CNs, which were annotated according to their mean cell type compositions (**Supplementary Table 2**).

For the end-to-end virtual MIF workflow, the same procedure was performed independently using only virtual stains. Cells were segmented from virtual DAPI using Mesmer, and virtual marker expression was quantified within the resulting virtual cell masks and independently normalized to [0,1]. Virtual cell types were defined by k-means clustering of the normalized virtual MIF profiles and annotated from their marker-enrichment profiles using the same centroid z-score procedure. The resulting virtual cell type definitions, together with their post hoc mapping to the unified biological annotations used for visualization, are provided in **Supplementary Data 2**. For each virtually segmented cell, a local composition vector was constructed from the cell types of its 10 nearest spatial neighbors, and these vectors were independently clustered to define virtual MIF CNs. The resulting CN compositions and annotations, together with the corresponding unified CN annotations used for case visualization, are provided in **Supplementary Data 3**. The measured and virtual workflows were therefore completed independently, without sharing cell masks, marker expression profiles, cell type assignments, or CN assignments. Unified cell type and CN annotations were assigned only after completion of both analyses and were used for visualization and biological comparison rather than for clustering or CN construction.

Supervised CN prediction was additionally evaluated by directly predicting the fixed real MIF-derived CN labels from virtual MIF or H&E without intermediate cell-type prediction. For these controlled comparisons, cells and spatial coordinates were defined using the real DAPI segmentation so that all models were evaluated on the same cells and CN labels. Models were evaluated using 5-fold grouped out-of-fold (OOF) prediction. For virtual MIF, the input comprised the normalized virtual marker expression vector of the central cell concatenated with the mean virtual marker expression vector of its 10 nearest neighboring cells. For H&E, a UNI2-h representation of the central cell was concatenated with the mean UNI2-h representation of its 10 nearest neighboring cells. Both approaches used a multilayer perceptron (MLP) with an output corresponding to the six esophagus or seven lung CN classes.

#### Patient-level downstream tasks

For patient-level analyses, we adopted shared markers across all virtual staining models, including DAPI, CD3e, CD8, CD20, CD68, Ki67, PDL1, and PanCK. Each MIF marker was encoded separately with a frozen DINOv3 image encoder. Each single-channel marker image was replicated along the channel dimension to form a three-channel input, which yielded a 768-dimensional DINOv3 embedding. The resulting marker features were combined using a learnable query pooling to form a patch-level MIF representation. H&E patches were encoded with the UNI2-h (R. J. Chen et al. 2024) pathology foundation model, using its 1,536-dimensional classification (CLS) token representation. Patch representations from a tissue region were aggregated into a single feature with an attention-based multiple instance learning (ABMIL, Ilse et al. 2018) model. Splits were made at the patient level to prevent patches from the same patient appearing in both training and evaluation partitions.

We first assessed whether virtual MIF generated from paired H&E images retained patient-level predictive utility comparable to that of real MIF across 4 H&E-MIF paired held-out cohorts. Survival prediction was optimized by minimizing the negative partial log-likelihood of the Cox proportional-hazards model (Katzman et al. 2018) and evaluated with the concordance index (C-index), whereas grade and classification tasks were optimized with cross-entropy loss and evaluated with AUROC.

We then evaluated 6 tasks in TCGA cohorts, for which real MIF was not available. For each task, virtual MIF alone and H&E combined with virtual MIF were compared with an H&E-only UNI2-h baseline. Classification tasks were evaluated by AUROC, and survival prediction tasks were evaluated by C-index as described above. The combined H&E and virtual MIF results were obtained by late fusion of the independently trained patient-level models. For each held-out patient, the H&E model and the corresponding virtual MIF model produced out-of-fold output scores. Scores from each modality were standardized separately within each output dimension and then summed with equal weight.

Bars and error bars report the mean and standard deviation, respectively, across the five held-out folds. Statistical confidence for comparisons in the main figure was assessed separately from the fold-based error bars. Differences between combined H&E–MIF and H&E-only performance were calculated from the paired out-of-fold patient predictions using 5,000 patient-level bootstrap resamples. A two-sided 95% bootstrap confidence interval was constructed for the metric difference, and an improvement was considered significant when its lower bound exceeded zero.

#### Regional contribution and feature-associated survival analysis

To identify patches that influenced MIL survival predictions, we removed one patch at a time from each region bag and recomputed the predicted risk score while keeping the trained model fixed. Patch contribution was measured as the change from the risk score obtained with the complete bag to the score obtained after removing that patch. A positive contribution indicates that including the patch increased the predicted risk, whereas a negative contribution indicates that it decreased the predicted risk. This procedure used repeated forward inference, and the model was not retrained. This analysis was applied to the held-out metastatic colorectal cancer survival and to TCGA-COAD PFS cohorts. Virtual MIF signals used in these analyses were generated using QUEST-semantic.

Feature-associated survival analyses were performed in TCGA-COAD using PFS as the endpoint. Patient-level virtual MIF summaries were generated for DAPI, PanCK, CD68, CD8, and CD20, and the CD8/PanCK ratio was calculated as an interpretable measure of cytotoxic T-cell signal relative to epithelial-tumor signal. For Kaplan-Meier analysis, patients were divided into high- and low-value groups at the cohort median for each virtual marker and for the CD8/PanCK ratio. Differences between curves were evaluated with two-sided log-rank tests.

A multivariable Cox proportional-hazards model (Cox 1972) included the patient-level out-of-fold (OOF) virtual-MIF MIL score, generated for each patient by the cross-validation model for which that patient was held out, together with the CD8/PanCK ratio, age, sex, disease stage, and microsatellite-instability status. Effect estimates are presented as log hazard ratios with 95% confidence intervals. A confidence interval that does not cross zero indicates that the variable remains independently associated with PFS after adjustment for the other covariates. Positive log hazard ratios indicate increased progression risk, whereas negative values indicate a protective association.

#### Cross-modality retrieval evaluation

We evaluated whether an H&E query could retrieve a biologically similar MIF patch from a reference gallery. For each query, a virtual staining model first generated the requested MIF panel from H&E. A frozen Eva (Yufan Liu et al. 2025) spatial proteomics encoder then converted the virtual MIF into a query embedding. Candidate real MIF patches were encoded with the same Eva encoder, and the gallery was ranked by cosine similarity between the query and candidate embeddings. All patches originating from the query’s source region were excluded from its gallery, preventing retrieval of a spatially adjacent or duplicate field. The real MIF paired with the query H&E was retained as an evaluation reference and was not used to rank candidates.

Haiku (Cui et al. 2026) was evaluated as a specialized cross-modal retrieval baseline. For Haiku, the H&E query and real MIF gallery images were encoded with its released modality-specific encoders and projection heads, and the resulting embeddings were normalized, and gallery images were ranked by cosine similarity. Haiku therefore did not generate an intermediate virtual MIF image.

Patch-level expression retrieval similarity was evaluated on Mixed 10K. For each query, marker expression vectors were obtained from its paired MIF reference and from the top-ranked MIF candidate. Retrieval quality was quantified by the cosine similarity and Euclidean (L2) distance between these expression vectors. Expression similarity was calculated using a fixed marker panel shared across patches in Mixed 10K, including CD8, CD4, CD3e, CD68, FoxP3, PDL1, PD1, Ki67, CD31, and DAPI. Differences between the L2-distance distribution of QUEST-semantic and another method were assessed with a two-sided two-sample Kolmogorov-Smirnov test.

Cellular retrieval was evaluated in held-out gastrointestinal stromal tumor (GIST) and pancreas cancer (PC) cohorts with cell segmentation and cell type annotations. The GIST cohort contained 1,319 patches from 25 regions, and the PC cohort contained 2,514 patches from 29 regions. Retrieval was performed separately within each cohort, and the two candidate galleries were not pooled. For each patch, a cell composition vector was constructed from the fraction of cells assigned to each cell type, and the dominant cell type was defined as the type with the largest fraction. For each query, we calculated the PCC between the cell-composition vectors of the paired MIF and the top-ranked retrieved MIF patch. We also reported top-1 dominant cell type accuracy, defined as whether the dominant cell type in the retrieved patch matched that in the paired MIF. Precision@k was defined as the proportion of the top-k retrieved patches whose dominant cell type matched that of the paired reference. All metrics were first calculated separately within each cohort and then averaged across the GIST and PC cohorts.

#### Zero-shot marker prediction

We evaluated zero-shot marker prediction with QUEST-semantic by completely excluding 24 markers from both model training and validation. Two zero-shot QUEST variants were trained from scratch using only the remaining markers, and neither model contained learnable marker-identity embeddings.

The two variants differed only in the text prompts used to represent markers throughout both training and inference. In the function prompt variant, the marker was conditioned on a description of its biological role. In the drawing prompt variant, the marker was instead conditioned on a description of its expected staining morphology, subcellular localization, and spatial distribution. Marker descriptions were encoded into 3,072-dimensional embeddings using OpenAI’s text-embedding-3-large model (https://developers.openai.com/api/docs/models/text-embedding-3-large) and projected into the shared marker-query space. At inference, unseen markers were requested using the same prompt type on which the corresponding variant had been trained. Zero-shot performance was evaluated on a fixed set of 10,000 patches sampled from the held-out test set. For each of the 24 unseen markers, pixel-level PCC between the virtual MIF prediction and real MIF reference was calculated and averaged across patches to obtain the per-marker result.

#### 3D virtual MIF construction

To construct a three-dimensional virtual MIF volume, we used the c001_A prostate cancer biopsy from the nondestructive 3D pathology dataset (Xie et al. 2022). The dataset contains an optically cleared intact tissue volume imaged using TO-PRO-3 nuclear and eosin fluorescence channels. The fused volume had an in-plane resolution of approximately 0.88 μm/pixel, and every second optical plane was converted into pseudo-H&E images to reduce redundancy and computational cost while preserving tissue continuity. Each H&E plane was processed independently by QUEST-semantic using overlapping 224×224 pixel sliding windows with a stride of 112 pixels. The predicted planes were then stacked in their original order to form the 3D virtual MIF volume. We showcased the resulting reconstructions for DAPI, CD31, EpCAM, and aSMA. No inter-plane registration was required because the planes were acquired through continuous optical sampling of the same intact tissue.

#### Computational efficiency and marker-panel scalability

Inference scalability was evaluated by requesting panels of 10, 20, 50, 100, and 500 markers from the same H&E patch under a fixed hardware and software configuration. We recorded inference time per patch and peak GPU memory allocated during inference. All measurements used a batch size of 1, and inference time for each model and panel size was reported as the median wall-clock time over 10 runs. For ROSIE, inference time included all forward passes required by its native sliding window procedure to cover the input patch. For fixed marker panel models, configurations exceeding the native panel size were constructed by expanding the final output layer to the requested number of channels with randomly initialized weights while retaining all preceding released weights. Particularly, MIPHEI-ViT did not generate all markers in a panel at once, and the shared encoder and decoder were retained, while the released marker-specific prediction heads were replaced with the requested number of randomly initialized heads. A 24-GB line corresponding to the memory capacity of a single NVIDIA RTX 4090 was included to indicate whether a configuration fit on a consumer-grade GPU. These measurements assess computational scalability only and were not included in the fidelity rankings.

### Implementation of related models

To benchmark virtual staining performance across datasets, we compared our models with recent methods applicable to H&E-to-MIF translation, including ROSIE, MIPHEI-ViT, GigaTIME, GigaTIME-flash, and Eva. Each H&E patch was resized to the spatial input dimensions required by the respective model. The predicted MIF maps were then resized back to 224×224 patch before evaluation. For cross-modal retrieval, we additionally compared against Haiku, a model specifically designed for this task. DINOv3 was used as the MIF feature extractor for downstream patient-level analyses. For all comparison methods, we used the publicly released weights provided by the original authors and followed their official inference procedures.

### ROSIE

ROSIE (Wu et al. 2025) uses a ConvNeXt (Z. Liu et al. 2022) backbone with a regression head to predict a 50-marker expression vector from each local H&E patch. Following the implementation, we applied the released model weights in a sliding window fashion across each input image and combined overlapping local predictions using a Gaussian weighting strategy, thereby converting the patch-level expression predictions into dense marker maps. The released weights of ROSIE can be found at https://huggingface.co/ericwu09/ROSIE.

### MIPHEI-ViT

MIPHEI-ViT (Balezo et al. 2026) combines a frozen H-optimus-0 (Saillard et al. 2024), ViT-g/14 pathology foundation-model backbone, LoRA adaptation, and a decoder to predict a fixed 16-marker MIF panel from H&E. The outputs of MIPHEI-ViT were mapped to the intensity range before evaluation following the released inference implementation. As MIPHEI-ViT was supervised on the ORION-CRC dataset, we treated it as an in-domain reference for that cohort rather than as a strictly zero-shot baseline. The released weights of MIPHEI-ViT can be found at https://huggingface.co/Estabousi/MIPHEI-vit.

### GigaTIME

GigaTIME (Valanarasu et al. 2026) is a UNet++-based (Zhou et al. 2020) model that translates H&E images into a fixed multiplex panel. Its original output is optimized for binary prediction, and we adapted the post-sigmoid outputs to evaluate the model’s pixel-level performance. The resulting probability maps were used throughout the same multi-scale benchmark as the other virtual-staining models. The released weights of GigaTIME can be found at https://huggingface.co/prov-gigatime/GigaTIME.

### GigaTIME-flash

GigaTIME-flash (Usuyama et al. 2026) is the efficient variant of GigaTIME, using a LoRA-adapted DINOv2 (Oquab et al. 2023) ViT-Small encoder and a convolutional decoder in place of the UNet++ architecture. We loaded the released weights and used the native marker positivity outputs without additional intensity matching. It shares the same marker panel and channel ordering as GigaTIME. The released weights of GigaTIME-flash can be found at https://huggingface.co/prov-gigatime/gigatime-flash.

### Eva

Eva (Yufan Liu et al. 2025) was used in two distinct roles, which were not assigned separate display names in the preceding text and figures but can be distinguished by task. The pretrained masked autoencoder checkpoint “Eva_model.ckpt” was used as the MIF feature encoder in the cross-modality retrieval task, which produced a 768-dimensional MIF embedding. The embeddings were used to construct the query and gallery representation. For virtual staining, we instead used the released H&E-to-MIF fine-tuned checkpoint “Eva_ft.ckpt”. In this mode, the marker names and H&E channels were provided as observed inputs, and the model reconstructed continuous intensity maps for the requested markers. The released weights of Eva can be found at https://huggingface.co/yandrewl/Eva.

### DINOv3

For patient-level downstream analyses, real and virtual MIF maps were encoded with a frozen DINOv3 (Siméoni et al. 2025) ViT-Base/16 model. Each single-channel marker image was replicated along the channel dimension to form a three-channel input compatible with the model’s input format. DINOv3 produced one 768-dimensional feature vector per marker and patch. Patch-level marker features were then aggregated and fed into the MIL models to obtain patient-level features for downstream tasks. We applied the released weights of DINOv3 built in PyTorch Image Models (timm) at https://huggingface.co/timm.

### UNI2-h

UNI (R. J. Chen et al. 2024) is a general-purpose pathology foundation model developed using the DINOv2 self-supervised learning framework. For the H&E-only reference in applicable patient-level tasks, we used its second-generation variant, UNI2-h, a ViT-H/14 model pretrained on more than 350,000 H&E and IHC slides. Each H&E patch was represented by the 1,536-dimensional class-token embedding returned by the frozen encoder, and these patch-level features were subsequently used for case-level prediction. We used the publicly released UNI2-h weights available at https://huggingface.co/MahmoodLab/UNI2-h.

### Haiku

Haiku (Cui et al. 2026) is a pretrained tri-modal representation model that aligns H&E, MIF, and clinical text inputs in a shared embedding space. Each query H&E patch was encoded directly with the Haiku H&E branch, while candidate MIF patches were encoded using the Haiku MIF branch and the corresponding marker names. The resulting H&E query and MIF gallery embeddings were used to calculate cosine similarity for retrieval. The released weights of Haiku can be found at https://huggingface.co/zhihuanglab/Haiku.

## Data and code availability

The public data can be found at their corresponding publications:

- TCGA: https://www.cancer.gov/ccg/research/genome-sequencing/tcga;
- PathoCell: https://huggingface.co/datasets/Kainmueller-Lab/PathoCell;
- ORION-CRC: https://zenodo.org/records/7637988;
- HEMIT: https://github.com/BianChang/HEMIT-DATASET;
- HuBMAP-intestine: https://portal.hubmapconsortium.org;
- 3D pathology: https://www.cancerimagingarchive.net/collection/pca_bx_3dpathology/.

The codebase, model checkpoints, and datasets are available at the following links:

- Codebase: https://github.com/YAndrewL/QUEST;
- Checkpoints: https://huggingface.co/yandrewl/QUEST;
- Tutorial datasets: https://huggingface.co/datasets/yandrewl/QUEST-tutorial-data;
- Held-out evaluation datasets: https://huggingface.co/datasets/yandrewl/QUEST-data.

## Acknowledgement

This work was supported by the University Research Committee of the University of Hong Kong and the Research Grants Council of Hong Kong, China, under the Early Career Scheme (27202625).

We extend our sincere gratitude to the developers of the ROSIE, GigaTIME, GigaTIME-flash, MIPHEI-ViT, DINOv3, UNI2-h, and Haiku models. Their commitment to the development and public release of their models has profoundly benefited the research community. These open-source models established the rigorous comparative standards that were instrumental in guiding the evaluation and development of this work.

## Author Contributions

Y.L. and O.F. developed the computational models and performed data analysis. O.F. and Y.D. contributed computational resources for model training and evaluation. Z.Z., L.Y., R.L., C.J., H.C., P.C., A.T.M., and A.E.T. facilitated data generation and offered advice and assistance for the project. Y.L. and Z.W. conceptualized the study and designed the methodology. Y.L., O.F., and Z.W. wrote the manuscript. All authors reviewed and edited the manuscript and approved its final version.

## Ethics, consent to participate, and consent to publish declarations

The use and computational analysis of the de-identified archival human tissue data in this study are approved by the Human Research Ethics Committee (HREC) of The University of Hong Kong. The specimens and associated data were retrospectively obtained by Enable Medicine from an existing biobank in accordance with applicable requirements for informed consent or waiver of consent, institutional review board or ethics committee oversight, and relevant regulatory guidelines. Given the retrospective nature of the study and the use of de-identified archival data, the requirement for informed consent to participate was waived by the HREC. Consent to publish was not applicable, as no identifiable individual participant information is reported.

