## Supplementary Figures and Tables for "Generalizable virtual staining from histopathology to spatial proteomics across biological scales"

#### Figures

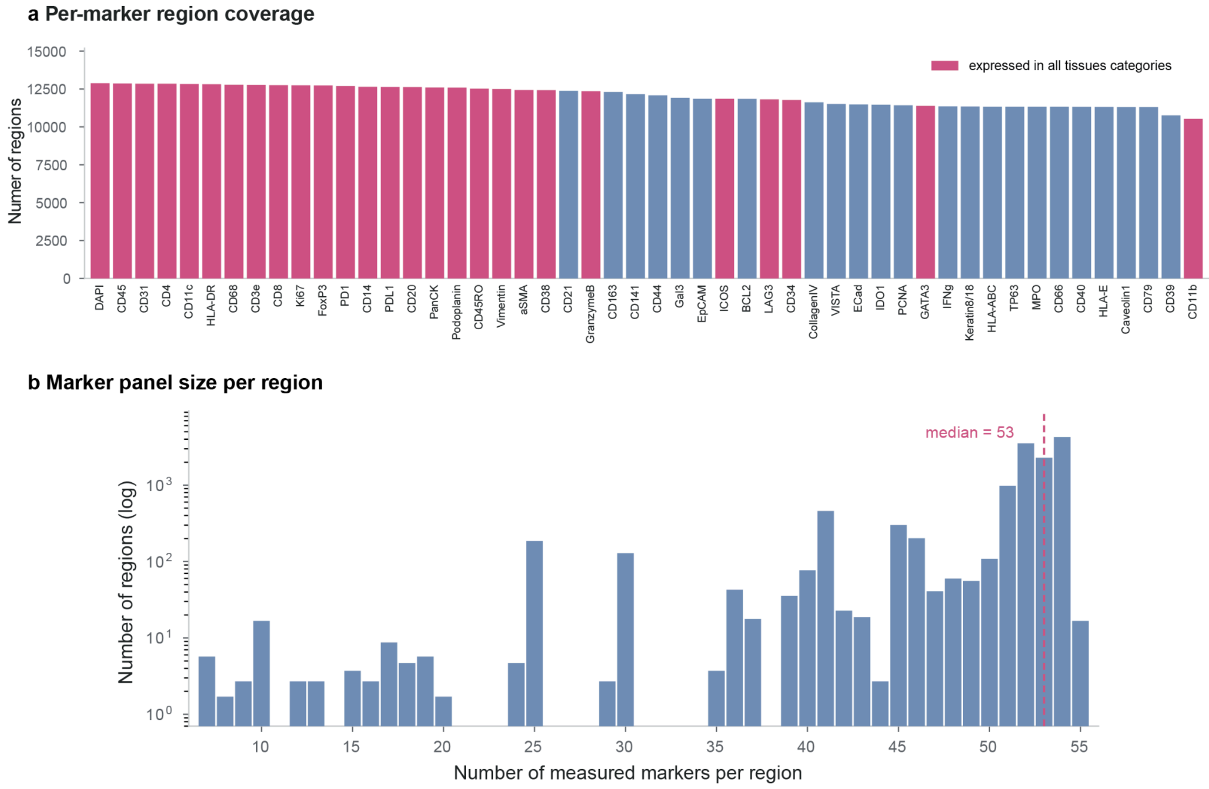


**Supplementary Figure 1: Marker coverage and panel size distribution across tissue regions. a,** Number of tissue regions in which each of the 50 most frequently measured markers was available. **b,** Distribution of marker-panel sizes across tissue regions, defined as the number of markers measured per region.

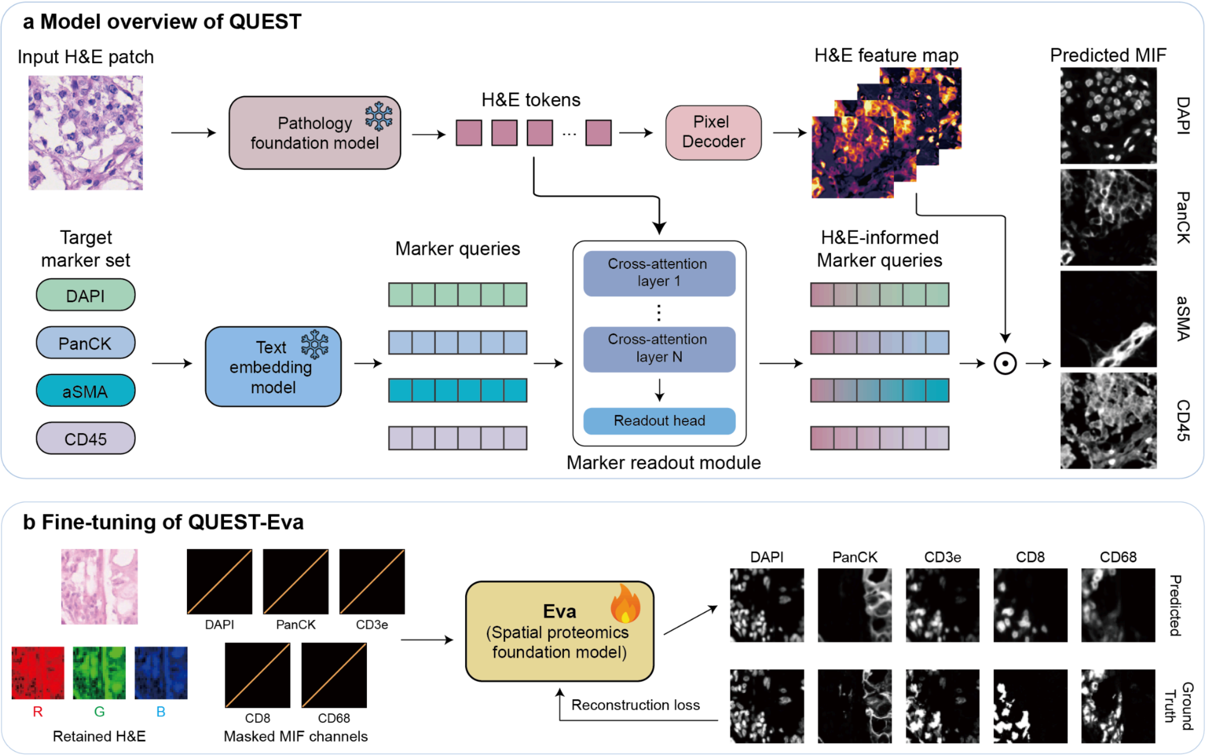


**Supplementary Figure 2: Model architectures of QUEST. a,** Overview of QUEST (**QUE**ry-based virtual **ST**aining). A frozen pathology foundation model encodes an input H&E patch into visual tokens, which are transformed by a pixel decoder into multi-scale H&E feature maps. Within the pixel decoder, a frozen ConvNeXt-Base branch extracts hierarchical convolutional features that are fused with the foundation-model tokens through a feature-pyramid network (**Methods**). In parallel, marker identity embeddings, or textual descriptions of the requested markers encoded by a frozen text-embedding model, are formed into marker queries. A marker-readout module uses stacked cross-attention layers to condition these queries on the H&E tokens. The resulting H&E-informed marker queries are applied to the feature maps to predict one MIF channel for each requested marker. **b,** Fine-tuning of Eva for H&E-to-MIF virtual staining. The H&E channels are retained while the target MIF channels are masked, and Eva is fine-tuned to reconstruct the MIF channels by minimizing a reconstruction loss.

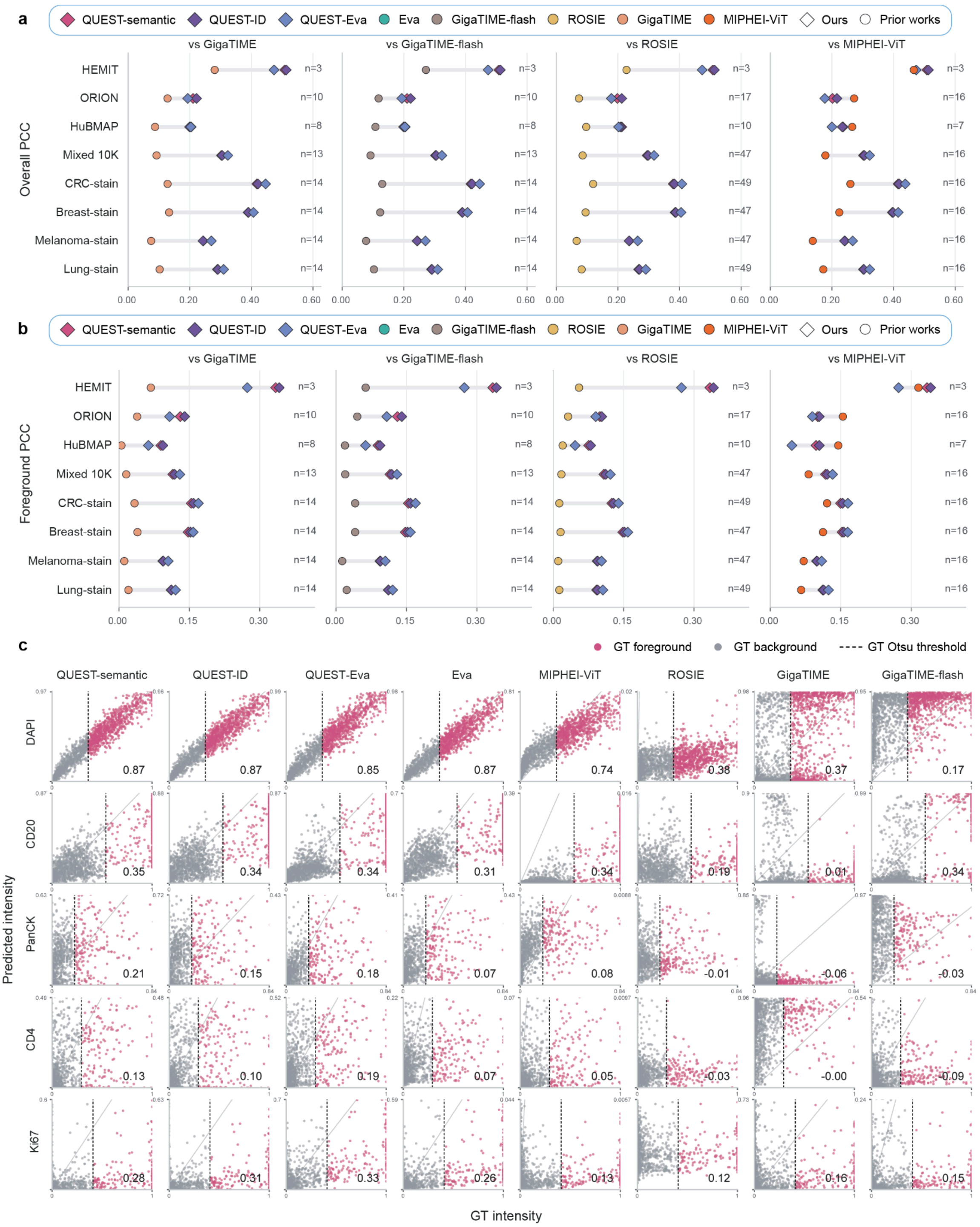


**Supplementary Figure 3: Overall and foreground PCC comparisons of common markers. a,** Pairwise comparisons of the pixel-level PCC across datasets. Each column compares the three models developed in this study with the indicated prior method. Within each dataset, evaluation was restricted to the marker set shared by all models included in that comparison. N denotes the number of these common markers. **b,** Corresponding comparisons of foreground PCC using the same model-specific common-marker sets. Foreground PCC was calculated using only pixels classified as foreground by an Otsu threshold applied to the real MIF signal. **c,** Scatter plots of predicted and real pixel intensities for representative markers. Pink and gray points denote ground truth foreground and background pixels, respectively.

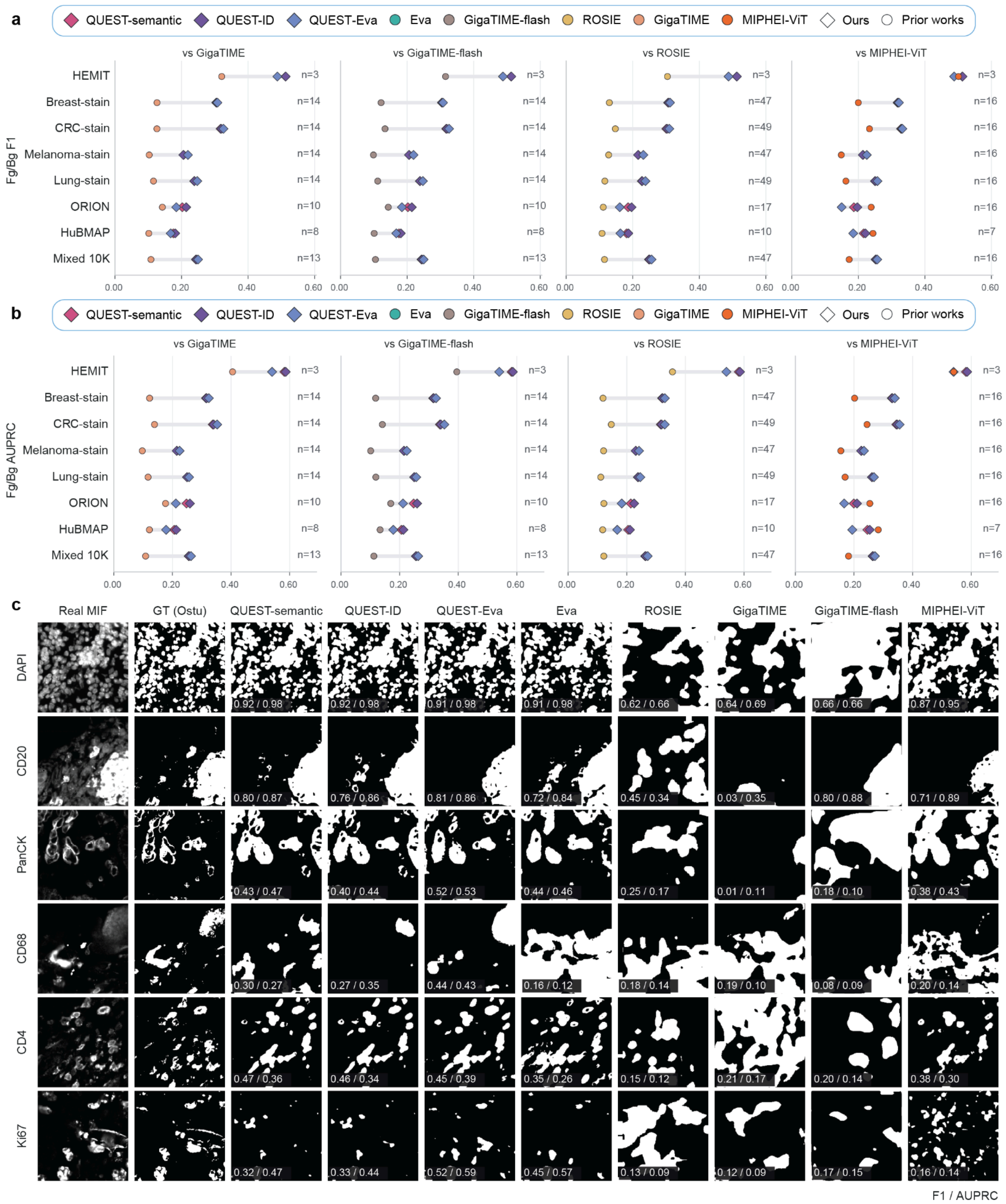


**Supplementary Figure 4: Foreground–background classification from continuous virtual staining predictions. a,** F1 scores for foreground–background classification across datasets. Each column compares the three models developed in this study with the indicated prior method. Within each dataset, evaluation was restricted to the marker set shared by all models included in that comparison. N denotes the number of these common markers. **b,** Comparisons using the area under the precision–recall curve (AUPRC) with the shared marker sets. Ground truth foreground pixels were defined by applying an Otsu threshold to the real MIF signal. **c,** Visual examples of real MIF images, Otsu-derived ground truth masks, and binarized predictions from each method. Values overlaid on each prediction report F1/AUPRC.

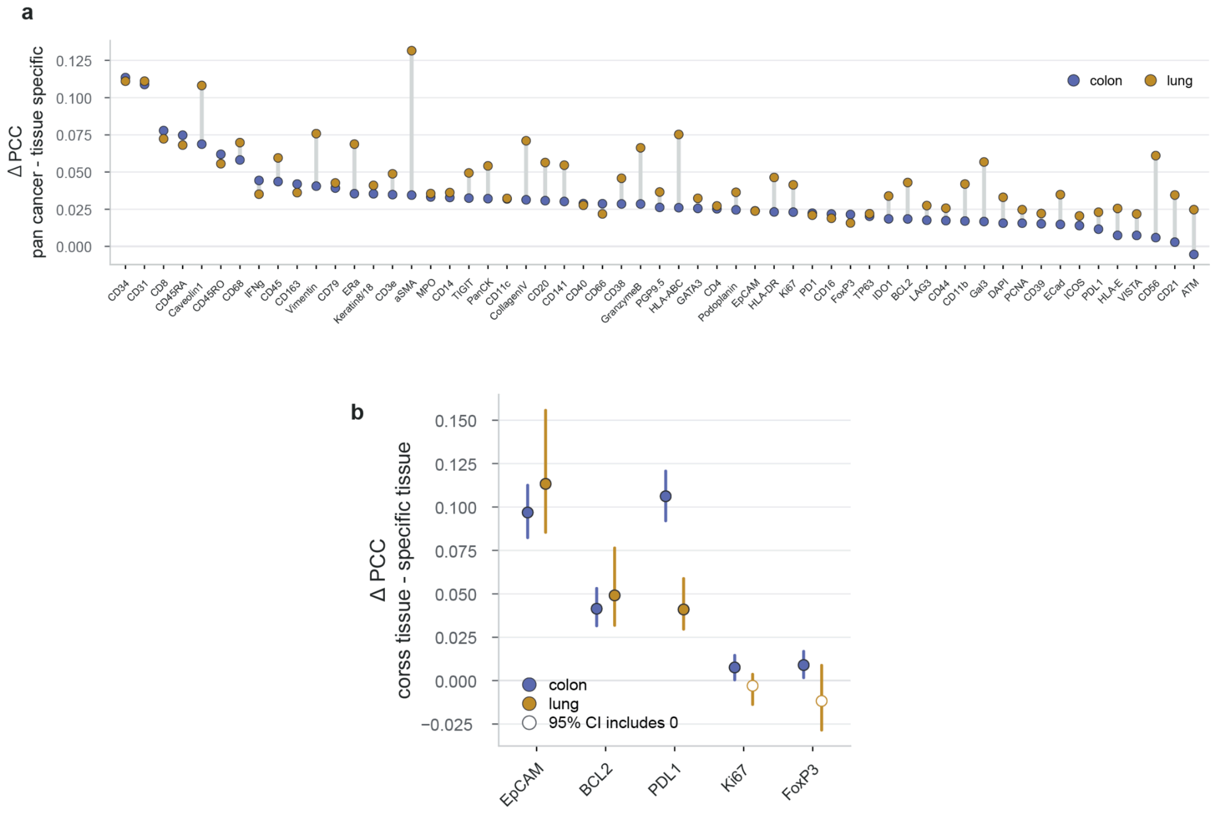


**Supplementary Figure 5: Pan-cancer training improves tissue-specific prediction and enables cross-tissue transfer of marker supervision. a,** Difference in per-marker PCC between the pan-cancer model and models trained exclusively on colon or lung data, evaluated on the corresponding held-out tissue test sets. Positive values indicate better performance of the pan-cancer model. The 57 markers evaluated in both tissues are shown and ordered by their mean improvement across colon and lung. **b,** Target tissue marker holdout analysis. Five markers were excluded from the target tissue during training (**Methods**). The difference in PCC denotes the discrepancy between the pan-cancer target tissue holdout model and the corresponding tissue-specific zero-shot model on held-out colon or lung data. Error bars indicate 95% confidence intervals, and open symbols denote intervals that include zero.

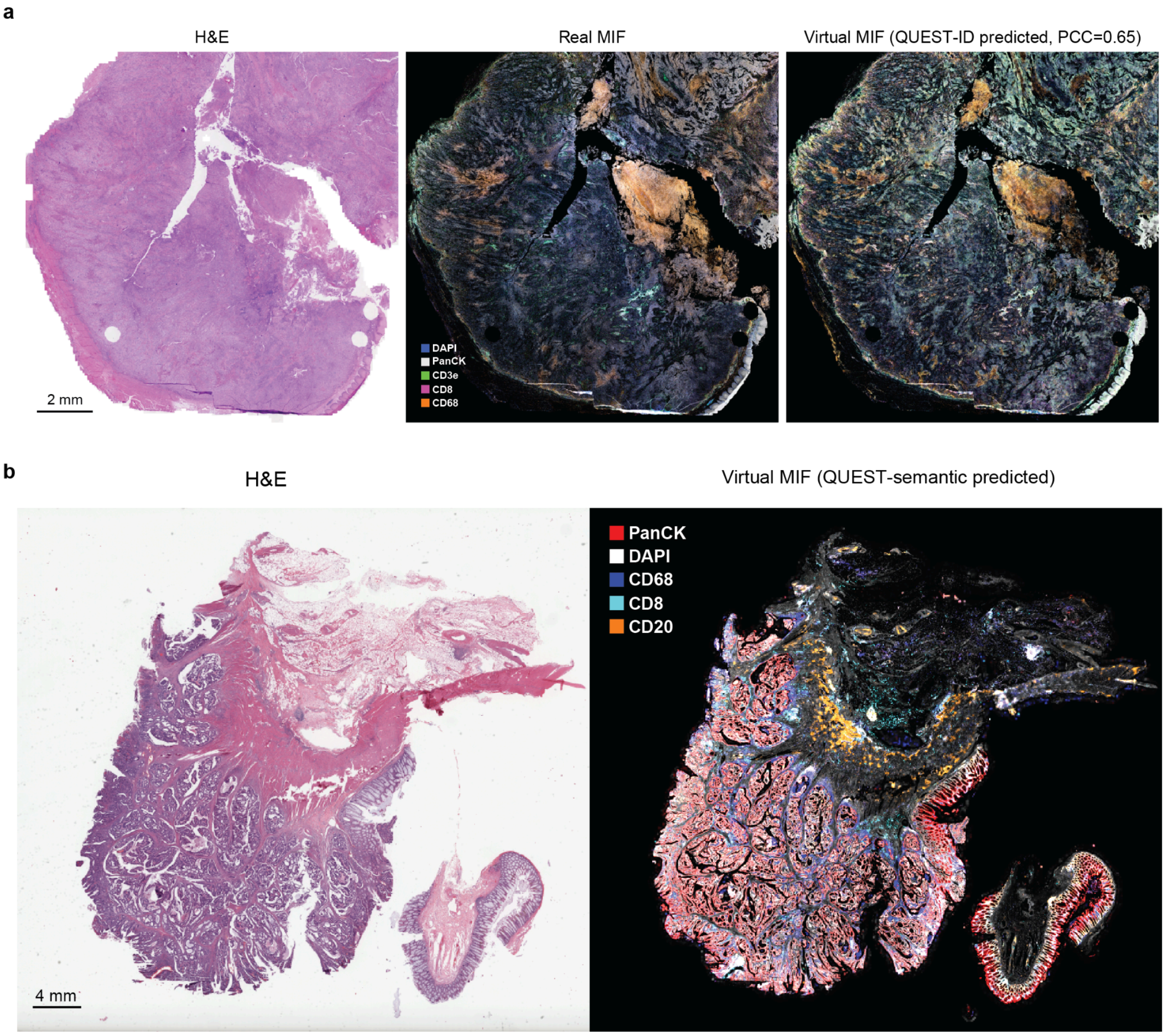


**Supplementary Figure 6: Virtual MIF prediction of whole slide images (WSI). a,** H&E, real MIF, and QUEST-ID-predicted MIF for a held-out WSI (PCC=0.65). **b**, H&E and QUEST-semantic-predicted MIF for an external TCGA WSI.

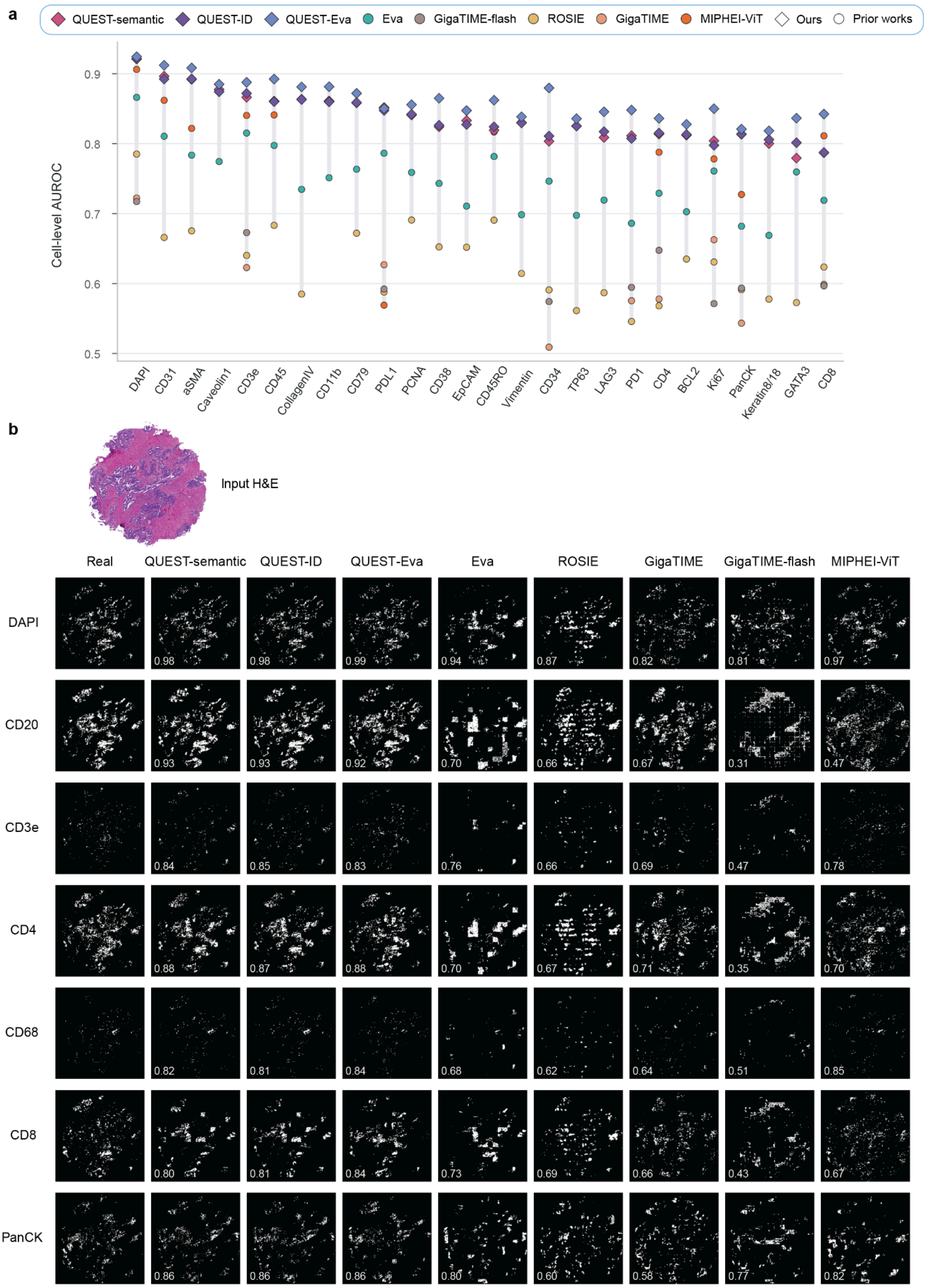


**Supplementary Figure 7: Binary cell-level evaluation of virtual MIF. a,** Per-marker cell-level AUROC for recovering marker-positive and marker-negative cell labels from continuous virtual MIF signals. **b,** Examples of binary cell-level maps for a held-out tissue region. For each marker, a threshold was applied to the real MIF signal to define the ground truth positive cells, yielding K positive cells. The real MIF column shows these positive cells, whereas each model column shows the K cells with the highest predicted signals. White therefore denotes ground truth positive cells in the real MIF column and the matched number of highest-ranked cells in each prediction, and black denotes all remaining cells or background. Values overlaid on the predicted maps report cell-level AUROC for the displayed marker and region.

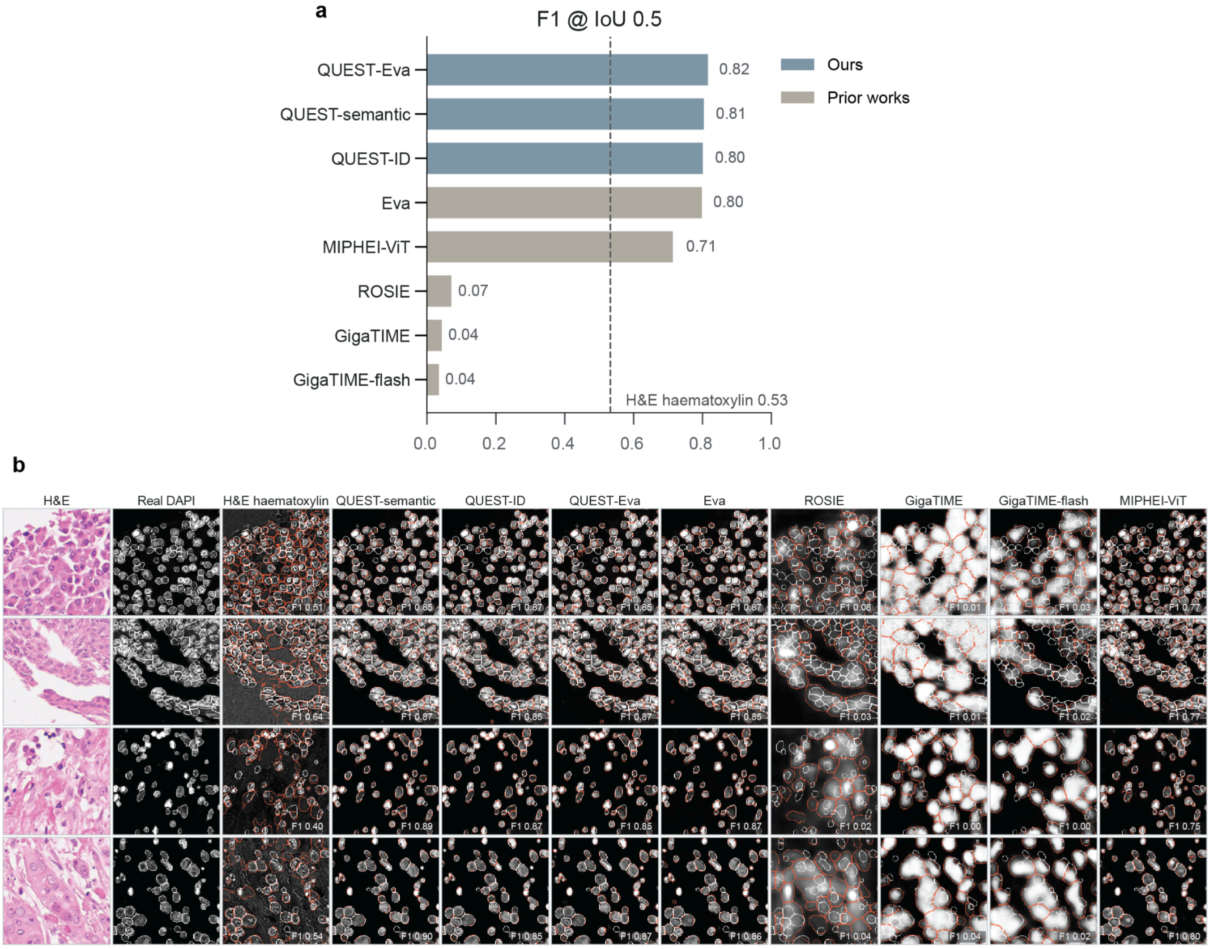


**Supplementary Figure 8: DAPI-based cell segmentation from virtual staining. a,** F1 scores for cell instances segmented from virtual DAPI images generated by each method. An intersection over union (IoU) of at least 0.5 is considered a true-positive match. The dashed line indicates the corresponding F1 score obtained from the hematoxylin channel extracted from the H&E images. **b,** Example H&E patches, paired real DAPI images, H&E-derived hematoxylin channels, and virtual DAPI predictions. White contours indicate reference cell instances derived from real DAPI, whereas orange contours indicate cell instances derived from the corresponding hematoxylin or virtual DAPI image. Values indicate the per-patch F1 score at IoU ≥ 0.5.

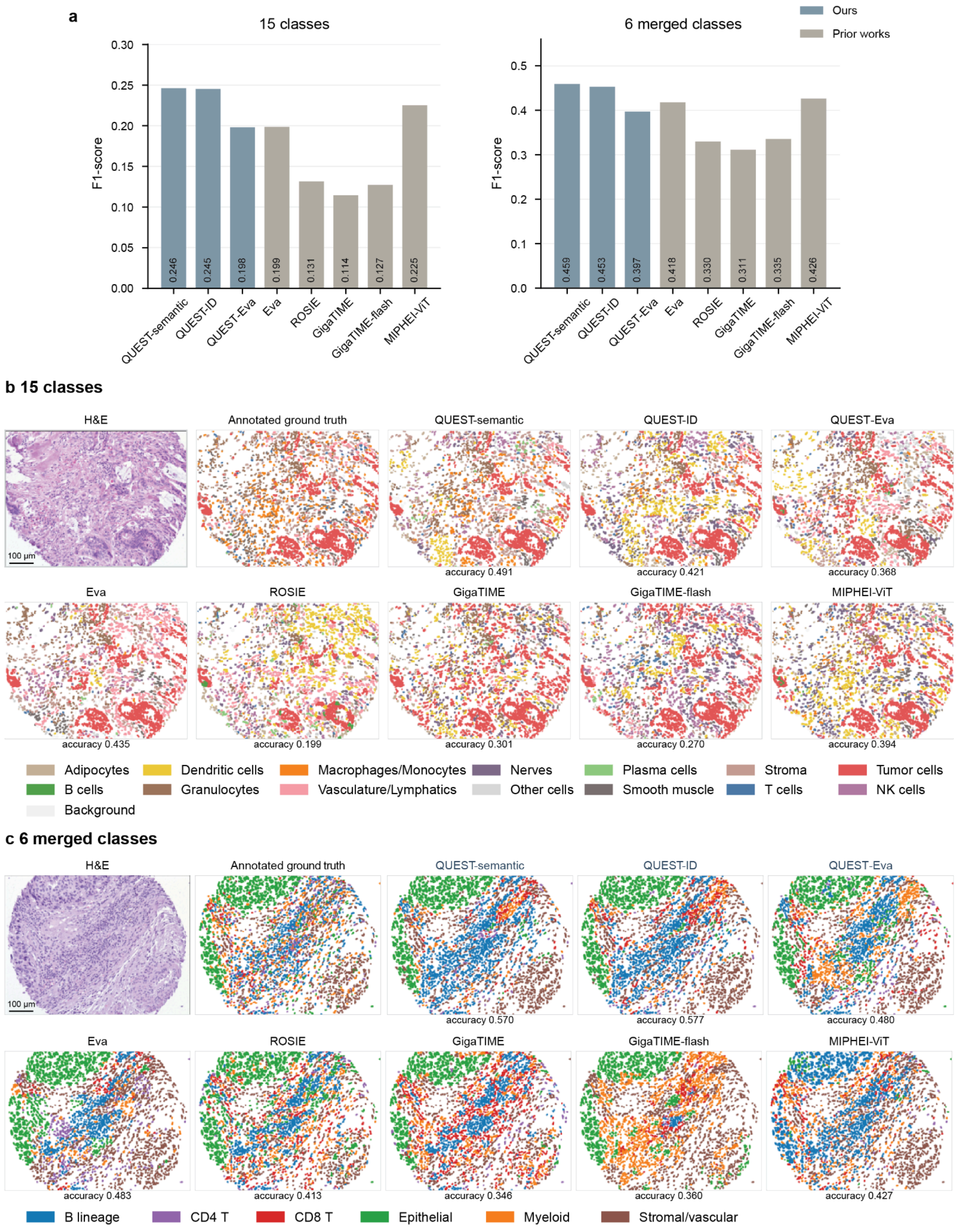


**Supplementary Figure 9: Cell typing from virtual MIF. a,** F1 scores for cell typing using the PathoCell's 15 cell classes and 6 merged cell classes. **b,** Representative cell-type maps for the 15 classes in an example tissue region. The paired H&E image and real MIF-derived annotations are shown alongside cell types inferred from each virtual MIF model. Values beneath the predicted maps report cell-level accuracy for the displayed region. **c,** Cell type maps after mapping cell types into 6 major classes: B lineage, CD4 T cells, CD8 T cells, epithelial cells, myeloid cells, and stromal/vascular cells.

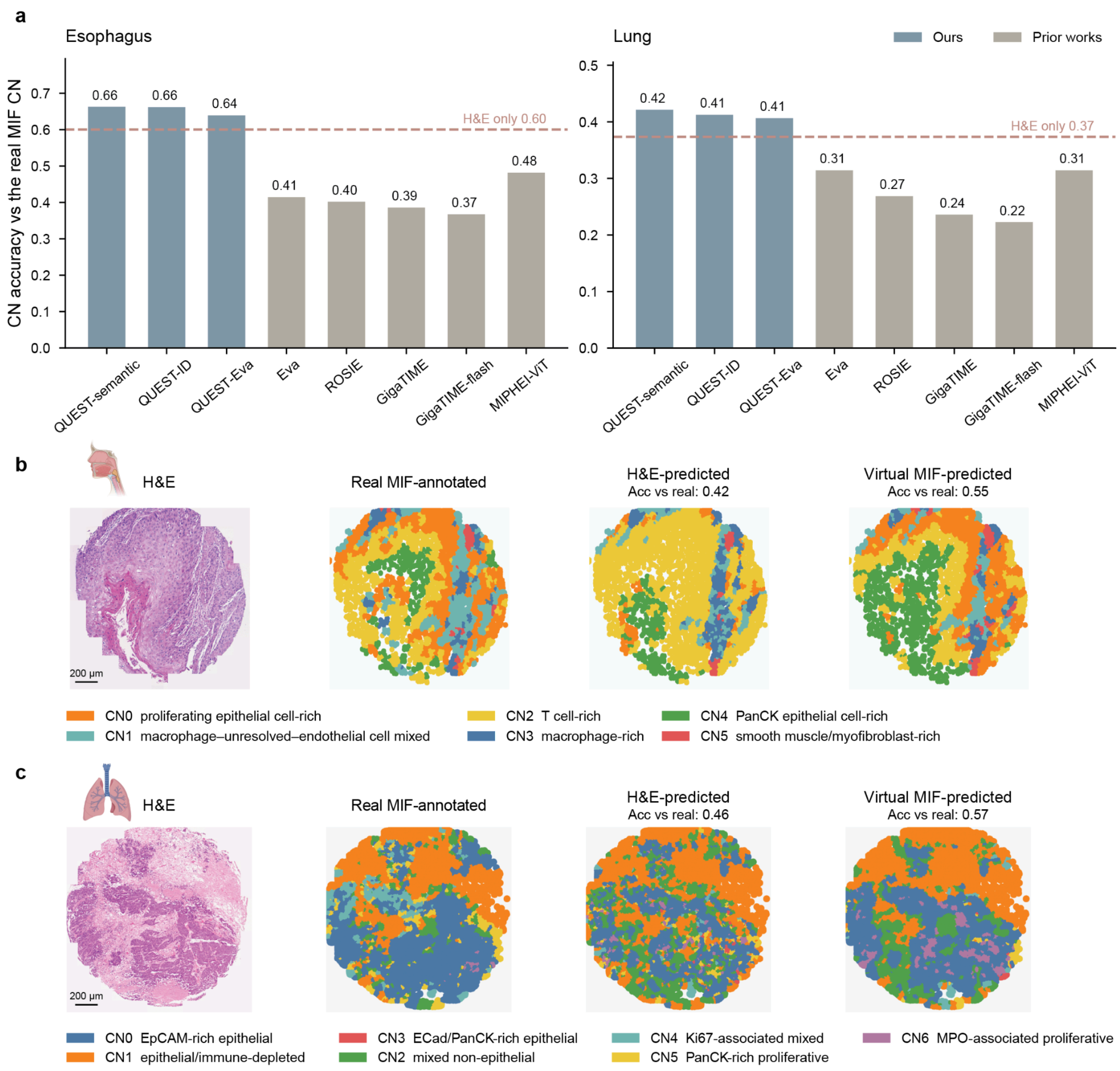


**Supplementary Figure 10: Virtual MIF supports cellular neighborhood annotation. a,** Cellular neighborhood (CN) accuracy relative to the real MIF reference in the esophagus and lung regions. All methods used the same cell masks derived from real DAPI. CN labels were predicted using supervised models trained on the corresponding virtual marker expression or H&E embedding. **b-c,** Representative CN maps for esophagus (**b**) and lung (**c**). Shown are CNs annotated from measured MIF, trained CN prediction from H&E, and trained CN prediction from virtual marker expression. All CN maps were visualized as Voronoi diagrams generated from cell centroids.

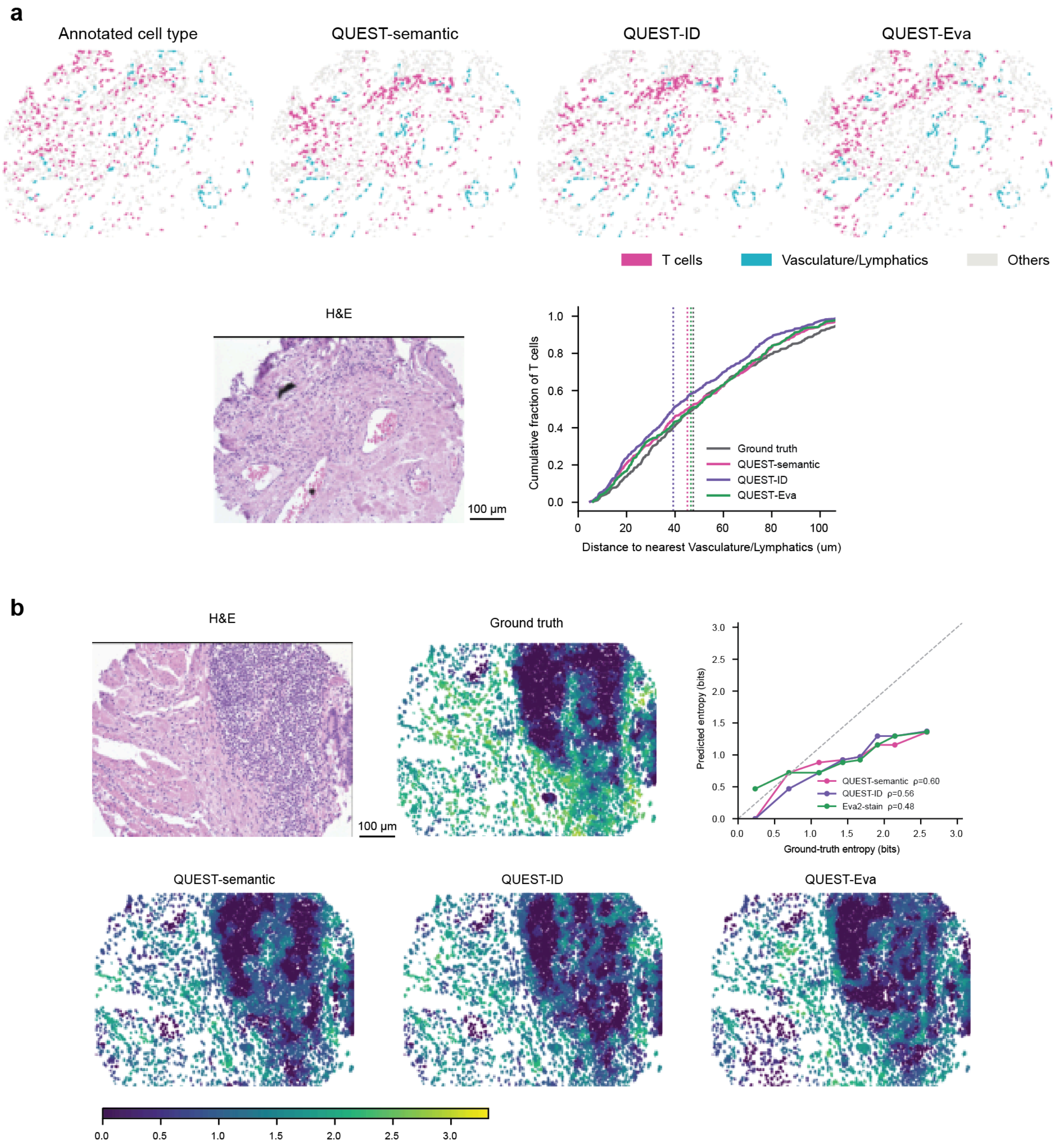


**Supplementary Figure 11: Virtual staining recapitulates cellular spatial organization. a,** Cell type maps derived from real MIF annotations and virtual MIF predictions from QUEST-semantic, QUEST-ID, and QUEST-Eva are shown for a representative tissue region. Cumulative distributions compare the distance from each T cell to its nearest vasculature/lymphatics cell in the annotated and predicted maps, and vertical dotted lines indicate the corresponding median distances. **b,** Local cell type diversity was quantified for each cell as the Shannon entropy of the cell type distribution among its 10 nearest neighbors. Maps show the ground truth entropy derived from real MIF and the corresponding entropy predicted from virtual MIF. Scatter plots compare predicted and ground-truth entropy, with the dashed diagonal indicating identity and Spearman's ρ reported for each model.

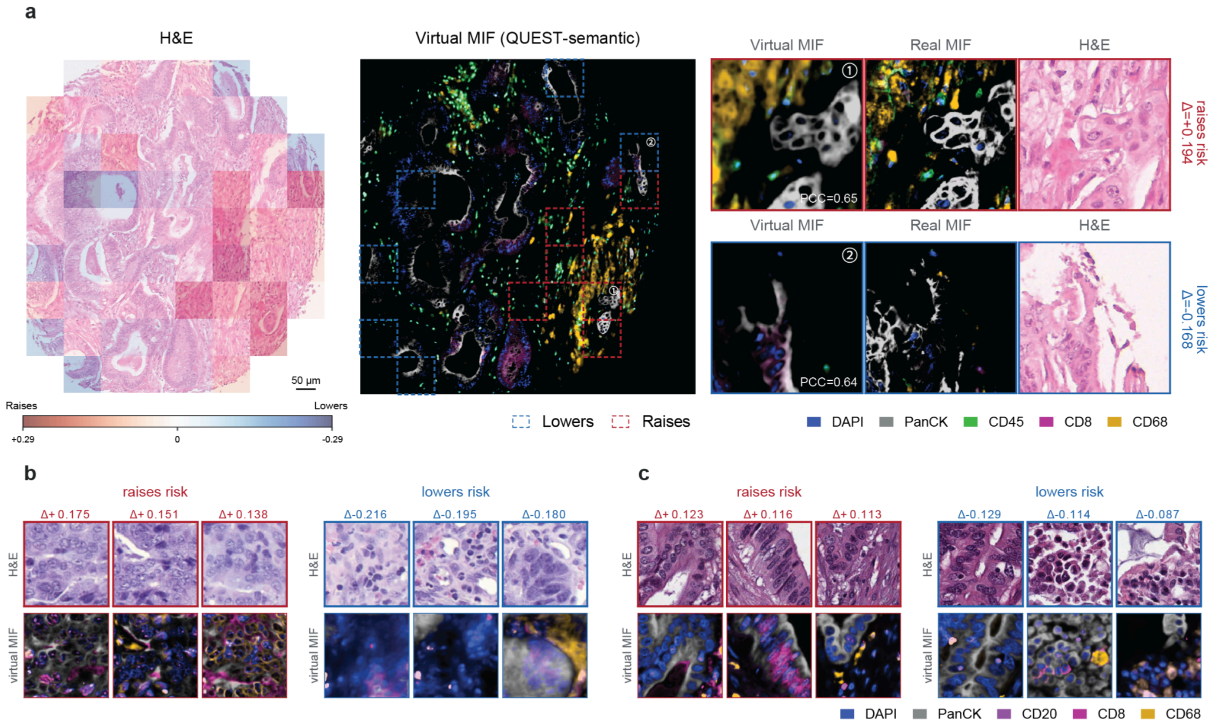


**Supplementary Figure 12: Patch-level attribution of clinical risk based on virtual MIF. a,** Patch-level contributions to a virtual MIF-based survival risk prediction in the metastatic CRC cohort. The contribution map highlights regions that increase or decrease predicted risk in red and blue, respectively. The corresponding whole-region virtual MIF is shown alongside zoomed-in examples of representative risk-raising and risk-lowering patches, with dashed boxes marking the top 10% of each. Zoomed-in views compare the virtual MIF, real MIF, and H&E images of risk-raising and risk-lowering patches. **b-c,** Spatial maps of virtual MIF-derived patch contributions to the predicted risk for two representative cases of TCGA-COAD cohort (**Methods**). Red and blue indicate patches that increase and decrease the predicted risk, respectively. For selected patches with positive or negative contributions, the corresponding H&E, virtual MIF images, and the contribution of predicted risk scores are shown.

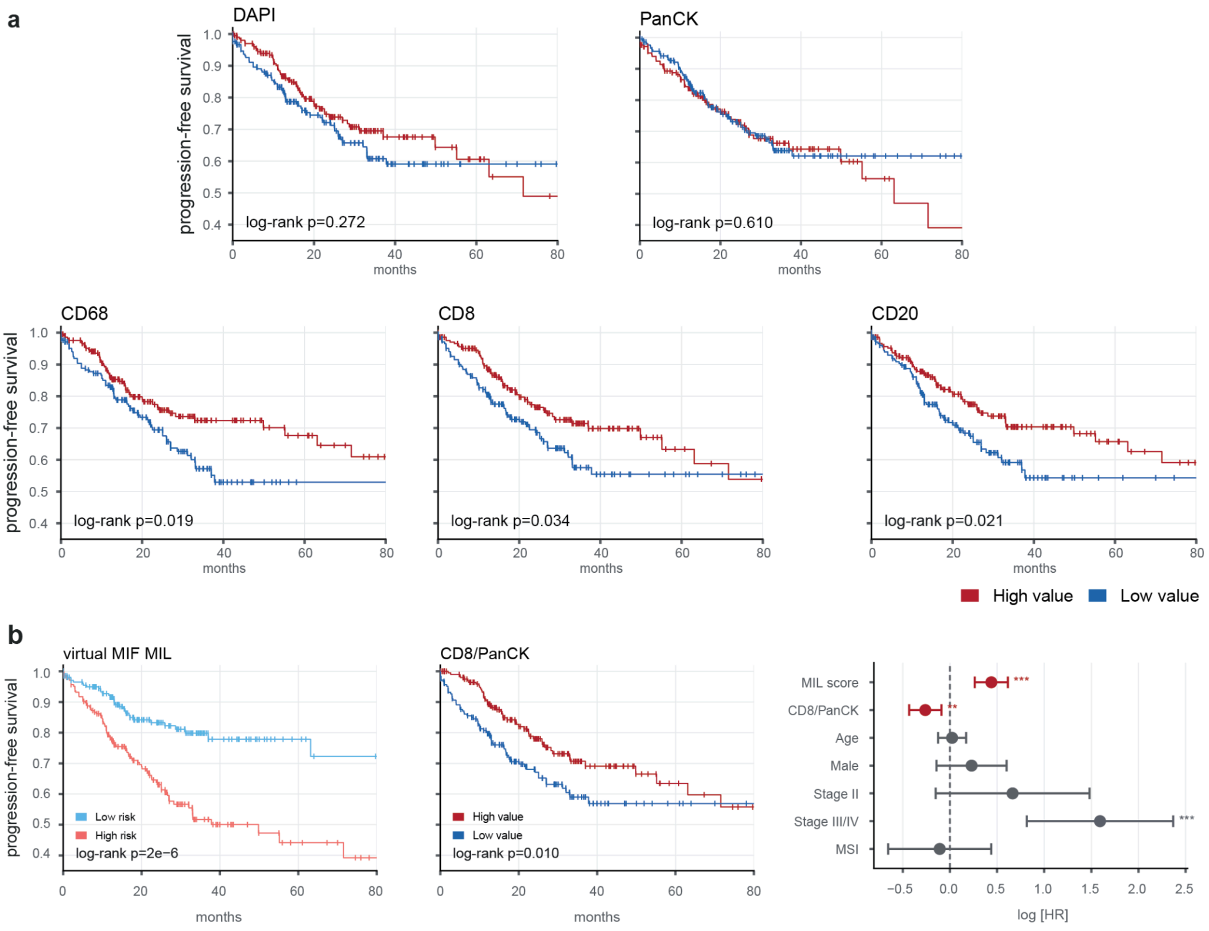


**Supplementary Figure 13: Progression-free survival analysis of virtual MIF-derived signals in TCGA-COAD. a,** Kaplan–Meier curves after stratifying patients into groups with high or low virtual MIF-derived DAPI, PanCK, CD68, CD8, and CD20 values. P-values were calculated using two-sided log-rank tests. **b,** Kaplan–Meier analyses after stratifying patients into high- and low-risk groups according to the virtual MIF-based multiple-instance learning (MIL) score (left) and into high- and low-value groups according to the virtual MIF-derived CD8/PanCK ratio (middle). The Cox proportional hazards analysis (right) includes the MIL score, CD8/PanCK ratio, age, sex, disease stage, and microsatellite instability status. Points and horizontal lines indicate log hazard ratios and 95% confidence intervals, respectively, and variables whose confidence intervals do not cross zero remain independently associated with progression risk after adjustment for the other covariates. Positive and negative log hazard ratios indicate increased and decreased progression risk, respectively. All analyses were performed in the TCGA colon adenocarcinoma cohort using progression-free survival as the endpoint (n=428).

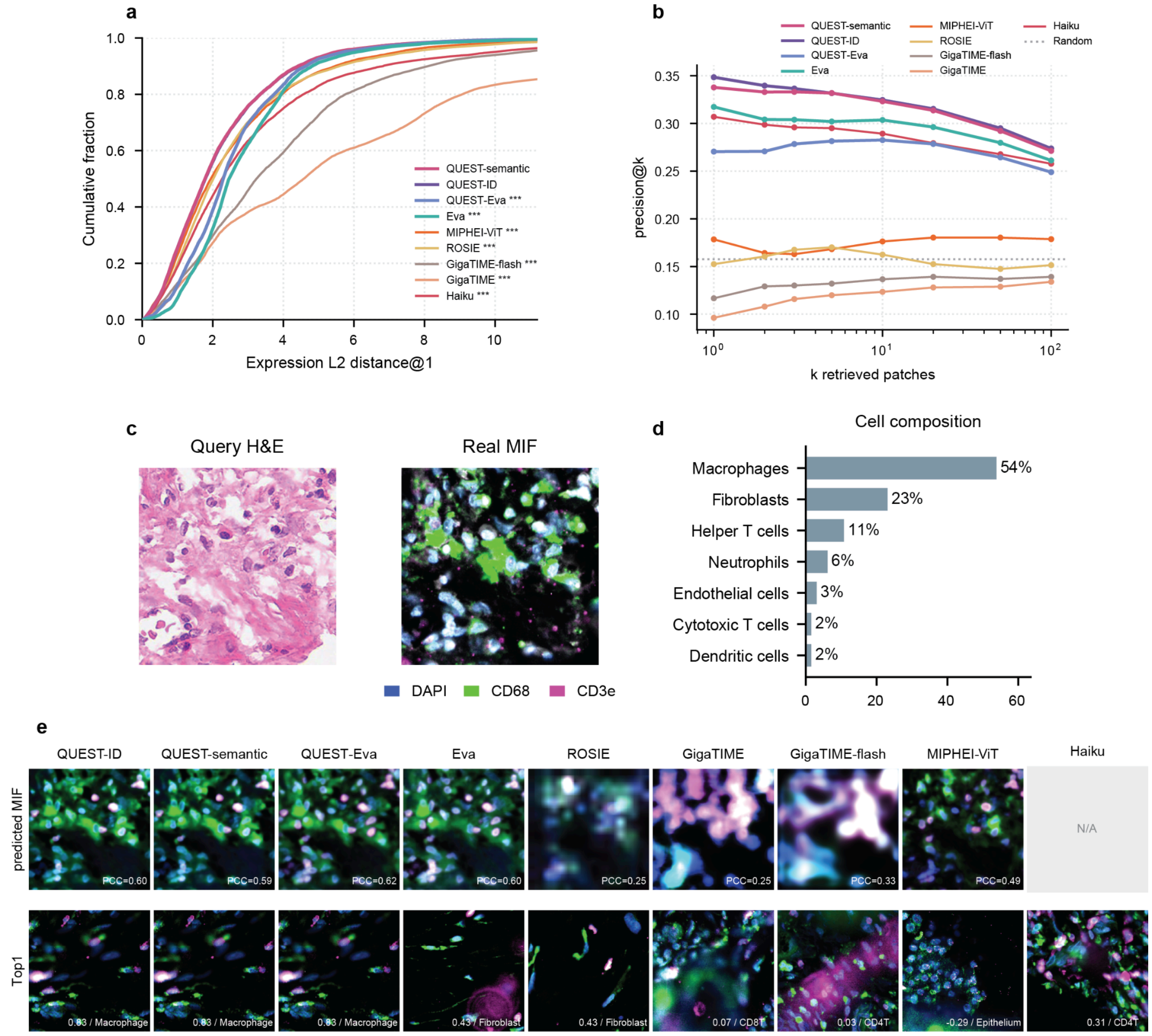


**Supplementary Figure 14: Evaluation of cross-modality retrieval performance. a,** Cumulative distributions of the L2 distance between the marker expression profiles of the paired real MIF query and the top-1 retrieved MIF patch. Statistical differences from QUEST-semantic were assessed using the Kolmogorov–Smirnov test (**Methods**). ***, $p < 0.001$. **b,** Precision@k for retrieving gallery patches whose dominant cell type matched that of the paired real MIF query. **c,** Example query H&E image and its paired MIF reference. **d,** Cell-type composition of the real MIF reference. **e,** Virtual MIF predictions and top-1 MIF patches retrieved by each method. Values overlaid on predicted MIF images report PCC with the paired real MIF. Values and labels on retrieved patches report cell composition PCC and the dominant cell type, respectively. Haiku directly performs retrieval and therefore does not produce a virtual MIF image.

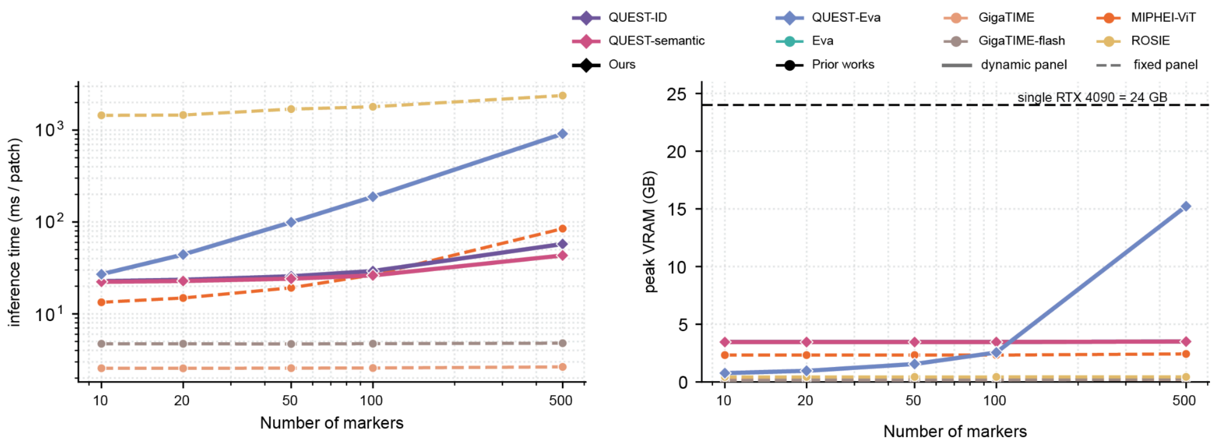


**Supplementary Figure 15: Computational efficiency and scalability with marker-panel size. a,** Inference time per H&E patch as the number of requested MIF markers increases from 10 to 500. **b,** Peak GPU memory consumption over the same range of marker-panel sizes. Solid lines indicate models that support dynamically specified marker panels, whereas dashed lines indicate fixed-panel methods. The black dashed line indicates the 24 GB memory capacity of a single consumer-grade NVIDIA RTX 4090 GPU.

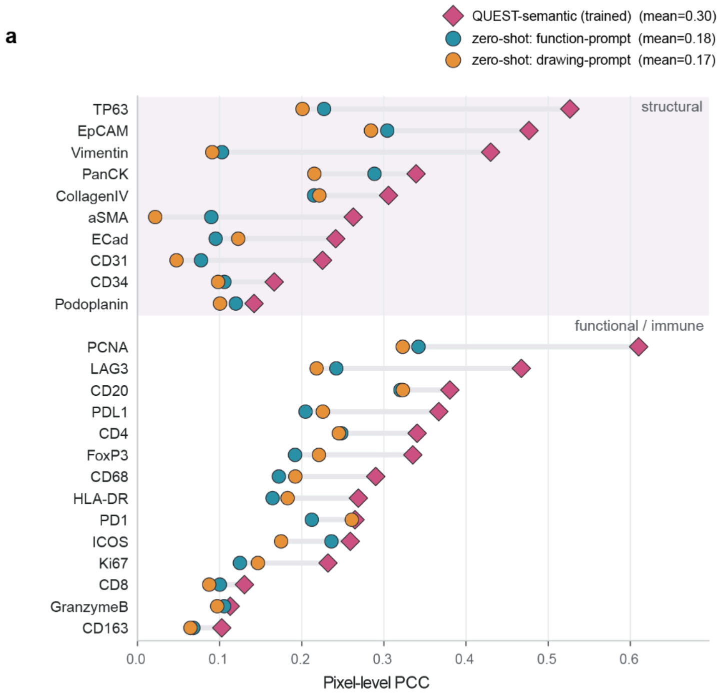


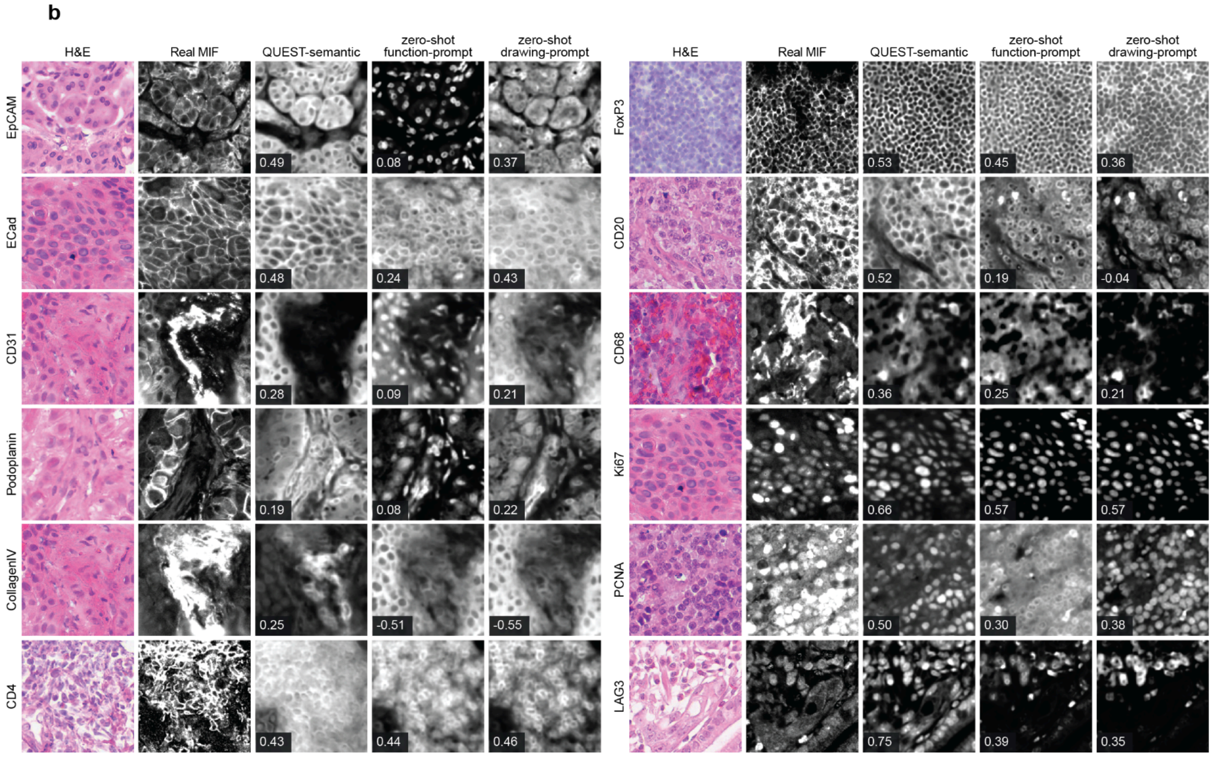


**Supplementary Figure 16: Zero-shot virtual staining of unseen markers using semantic prompts. a,** Pixel-level image PCC for 24 markers that were completely excluded from zero-shot model training (**Methods**), grouped into structural and functional/immune markers. Performance is shown for the supervised QUEST-semantic reference trained with these markers and two zero-shot variants conditioned on either function prompts describing the biological roles of the markers or drawing prompts describing their expected staining morphology and spatial localization. **b,** Visualizations of H&E inputs, real MIF references, and virtual MIF predictions for selected withheld markers. Values overlaid on predicted images report full-image PCC.

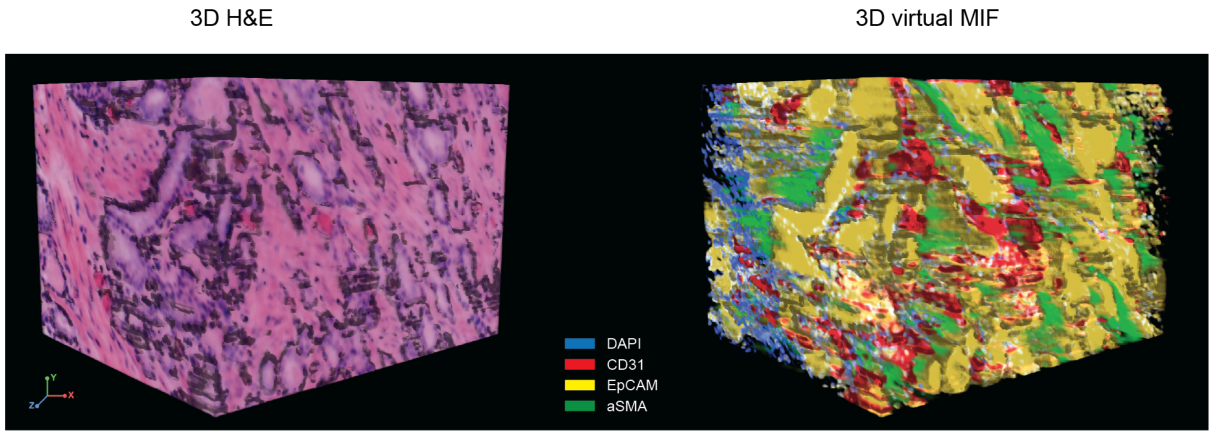


**Supplementary Figure 17: Three-dimensional virtual MIF reconstruction from H&E.** Volume renderings of a single prostate cancer sample are shown for the 3D H&E input (left) and the corresponding QUEST-generated virtual MIF reconstruction (right). Virtual marker predictions from the sampled optical planes were stacked to construct the 3D volume. Gradient-based surface shading was applied to the 3D volume renderings to enhance depth perception and did not alter the underlying signal intensities.

### Tables

**Supplementary Table 1: Marker enrichment profiles and cell type annotations for MIF-derived clusters.**

| Tissue | Enriched marker signals (z-score) | Cell-type annotation |
|---|---|---|
| Esophagus | CD45 (2.5), CD68 (2.4), HLA-DR (2.2) | macrophage |
| Esophagus | Ki67 (2.5), CD8 (2.3), EpCAM (2.2) | proliferating epithelial cell |
| Esophagus | CD3e (1.9), ECad (1.8) | T cell |
| Esophagus | PanCK (2.3), CD3e (1.3) | PanCK epithelial cell |
| Esophagus | aSMA (2.6) | smooth muscle/myofibroblast |
| Esophagus | None | unresolved |
| Esophagus | CD31 (2.6) | endothelial cell |
| Esophagus | FoxP3 (2.6), MPO (1.4) | FoxP3/MPO-associated cell |
| Lung | CD45 (2.0), CD4 (1.8), CD3e (1.1) | helper T cell |
| Lung | CD20 (1.9), FoxP3 (1.5), CD4 (1.3) | CD20/FoxP3-associated cell |
| Lung | None | unresolved |
| Lung | CD163 (3.0), Podoplanin (3.0), CD20 (2.7) | CD163/Podoplanin-associated cell |
| Lung | EpCAM (1.2) | epithelial cell |
| Lung | aSMA (3.6), Vimentin (1.0) | smooth muscle/myofibroblast |
| Lung | Podoplanin (1.8), CD163 (1.4) | Podoplanin/CD163-associated cell |
| Lung | ECad (3.3), PanCK (1.3) | ECad/PanCK epithelial cell |
| Lung | PanCK (2.4) | PanCK epithelial cell |
| Lung | CD8 (3.6), CD45 (1.9), CD3e (1.2) | cytotoxic T cell |
| Lung | CD68 (3.1), HLA-DR (1.6), Vimentin (1.5) | macrophage |
| Lung | CD31 (3.9), Vimentin (2.8), aSMA (1.1) | endothelial cell |
| Lung | Ki67 (2.2), MPO (1.8), EpCAM (1.1) | MPO-associated proliferating cell |
| Lung | Ki67 (2.9) | proliferating cell |

**Supplementary Table 2: Cell type compositions and annotations of MIF-derived cellular neighborhoods.**

| Tissue | Cell type composition | CN annotation |
|---|---|---|
| Esophagus | proliferating epithelial cell, macrophage | proliferating epithelial cell-rich |
| Esophagus | macrophage | macrophage-rich |
| Esophagus | macrophage, unresolved, endothelial cell, proliferating epithelial cell, smooth muscle/myofibroblast, FoxP3/MPO-associated cell | macrophage–unresolved–endothelial cell mixed |
| Esophagus | T cell, proliferating epithelial cell, PanCK epithelial cell, macrophage | T cell-rich |
| Esophagus | PanCK epithelial cell, T cell, macrophage | PanCK epithelial cell-rich |
| Esophagus | smooth muscle/myofibroblast, macrophage, unresolved, endothelial cell | smooth muscle/myofibroblast-rich |
| Lung | epithelial cell, CD20/FoxP3-associated cell, unresolved | EpCAM-rich epithelial |
| Lung | unresolved, PanCK epithelial cell | epithelial/immune-depleted |
| Lung | unresolved, helper T cell | mixed non-epithelial |
| Lung | ECad/PanCK epithelial cell, unresolved | ECad/PanCK-rich epithelial |
| Lung | unresolved, proliferating cell | Ki67-associated mixed |
| Lung | unresolved, PanCK epithelial cell, proliferating cell | PanCK-rich proliferative |
| Lung | MPO-associated proliferating cell, proliferating cell, epithelial cell | MPO-associated proliferative |
